# Aromatic Patch in WhiB-Like Transcription Factors Facilitates Primary Sigma Factor Interaction in *Mycobacterium tuberculosis*

**DOI:** 10.1101/2025.06.05.653497

**Authors:** Daisy Guiza Beltran, Tao Wan, Javier Seravalli, Magdaléna Horová, Camden Jones, Shanren Li, Pengchong Ma, Chloe Ong, Zhifang Lu, Donald F Becker, Jeffrey P. Mower, Qiuming Yao, Yu Pan, Hongfeng Yu, Adrie J. C. Steyn, LiMei Zhang

## Abstract

WhiB-like (Wbl) family proteins are a unique family of iron-sulfur ([4Fe-4S]) cluster-bound transcription factors found exclusively in Actinobacteria and actinobacteriophages, including the notoriously persistent pathogen *Mycobacterium tuberculosis* (*Mtb*). Despite their critical roles in cell development, stress response and antibiotic resistance, the mechanisms of gene regulation by the Wbl family proteins are not fully understood due to the lack of a canonical DNA-binding motif in most Wbl proteins. Here, we present structural and biochemical evidence demonstrating that all *Mtb* Wbl proteins bind to the same site in the conserved region 4 of the primary sigma 70 factor facilitated by a previously unrecognized structural motif, the aromatic patch, in the Wbl family. Our phylogenetic findings provide compelling evidence for a complex evolutionary relationship of Wbls between actinobacteria and the associated phages. Together, this work fills a critical gap in our understanding of the function, mechanism and evolutionary origin of Wbls.

## INTRODUCTION

The *WhiB-like* (*wbl*) proteins are a group of monomeric [4Fe-4S] cluster-containing transcription factors discovered in *Streptomyces coelicolor* (*Sco*) in 1972 and named after the white colonies of the *wbl* mutants due to the disruption of pigment formation (1). Subsequent genetic and bioinformatic analyses find that the Wbl proteins are widely distributed in the phylum Actinobacteria and their associated actinobacteriophages, but not in any other living organisms. Notably, several actinobacterial species of medical and industrial significance encode multiple Wbl paralogues (2–4). For example, seven Wbl members (WhiB1-7) are encoded in *Mtb*, 12 in *Sco* and four in *Corynebacterium glutamicum* (*Cglu*). Wbl proteins play critical roles in diverse biological processes in these actinobacteria, such as cell division and development, responses to redox stress and nutrition starvation, antibiotic resistance and pathogenesis (3,5–21). Because of that, the Wbls proteins have attracted a great deal of attention since their discovery (see recent reviews in (4,22)).

Wbl family proteins are characterized by two conserved motifs, the [4Fe-4S]-cluster binding motif composed of four conserved cysteines and the Gly-rich motif (also referred to as the β-turn) (Fig. 1A) (3,23). Based on their sequence similarity, these Wbl proteins are grouped into five major subfamilies, each represented by an *Mtb* Wbl (WhiB1-4, and WhiB7, respectively) and distributed across a wide range of Actinobacterial species (2,4). The rest of the Wbl proteins are more species-specific, such as WhiB5 and WhiB6 found only in Mycobacteria. Several subfamily-specific functional motifs have been reported, most notably the well-characterized “AT-hook” DNA binding motif in the WhiB7 subfamily (24–28). However, most Wbl proteins lack canonical DNA binding motifs, or no other Wbl family-wide motif apart from the two conserved motifs mentioned above has been reported. This knowledge gap has long posed a challenge in understanding how Wbl proteins regulate gene expression.

**Figure 1.**
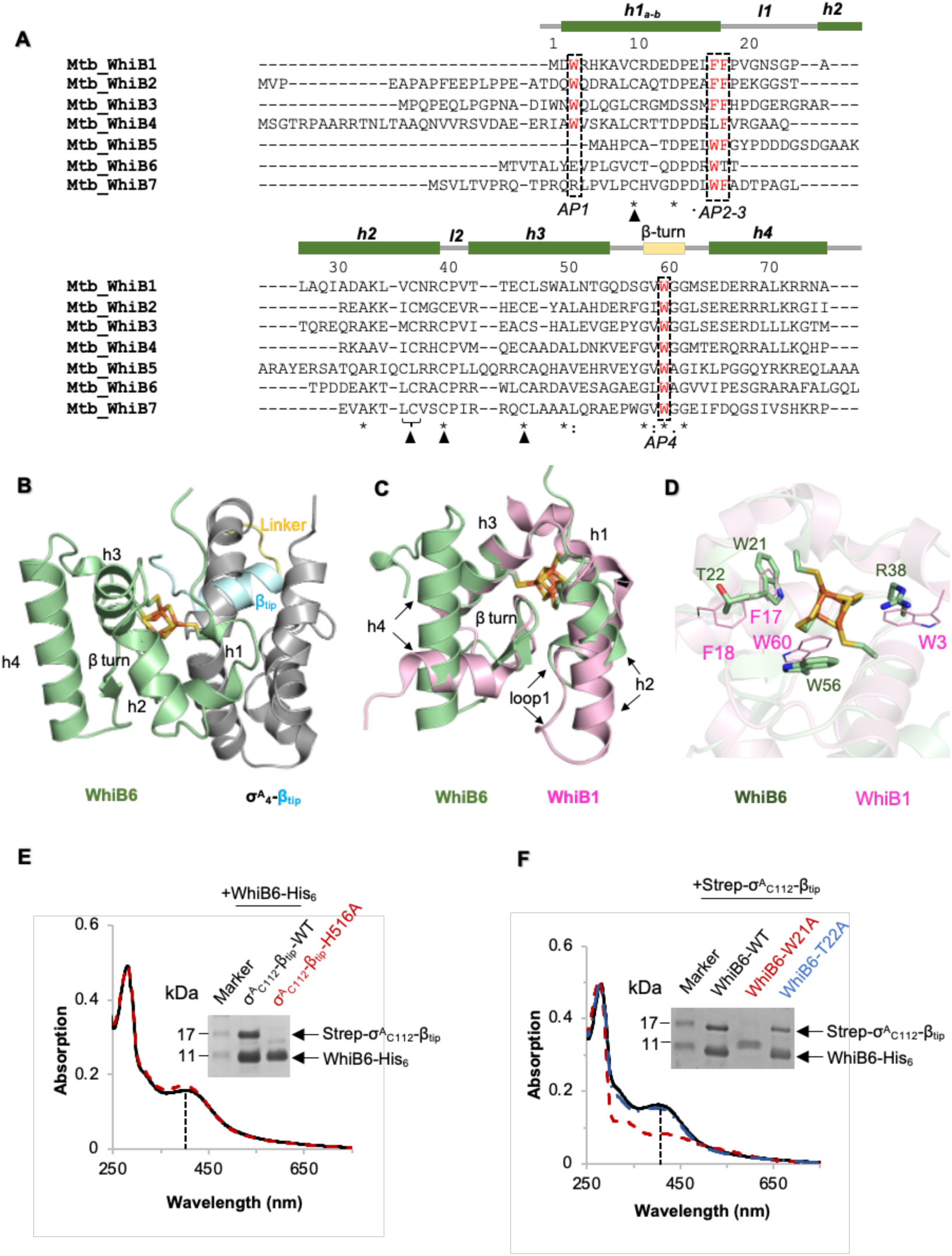
The aromatic patch (AP) facilitates *Mtb* WhiB6 binding to σ^A^_4_. (A) Sequence alignment of the seven *Mtb* Wbl proteins. The two previously identified motifs in the Wbl family, including the four conserved cysteines (the black triangle) in the [4Fe-4S] cluster binding motif and the Gly-rich β-turn, are highlighted. The assignment of the secondary structures (helices *h1-h4*, loop *l1-l2*, β-turn) is based on the σ^A^_4_-bound WhiB1 (PDB ID:6ONO). The residues in the aromatic patch (AP) in *Mtb* Wbls corresponding to W3, F17, F18, and W60 (referred to as AP1-4, respectively) in WhiB1 for σ^A^_4_ binding are highlighted in red fonts within the black frame. (B) Cartoon representations of the WhiB6:σ^A^_4_-β_tip_ complex. The key pair of residues, H516-P517, of σ^A^_4_ for WhiB6 binding is highlighted in sticks. (C-D) Structural overlay of σ^A^_4_-bound WhiB6 (pale green) with WhiB1 (pink, PDB ID:6ONO) and a close-up view of the Wbl:σ^A^_4_ interface around the Fe-S cluster binding pocket. The AP1-4 of WhiB1 and corresponding residues in WhiB6 are highlighted in sticks, together with the H516-P517 pair of σ^A^_4_ in the WhiB6:σ^A^_4_ complex. σ^A^_4_ and the [4Fe-4S] cluster of WhiB1 are not shown for clarity. In all structures, the helices *h1-4* and β-turn are indicated. (E and F) UV-visible spectra and SDS-PAGE analyses of the pull-down samples of WhiB6 and σ^A^_4_-β_tip_ (either wildtype [WT] or mutant as indicated). The color of the UV-visible spectra matches that of the sample labels. The intensity of the absorption peak around 410 nm, highlighted with dashed lines, is indicative of the occupancy of the [4Fe-4S] cluster in the purified samples. The absorption spectra were normalized at 280 nm. The uncropped SDS PAGE images are shown in Fig. S3.

The discovery that WhiB3 requires interaction with the conserved C-terminal region 4 of the primary σ^70^ sigma factor (σ^A^_4_) in the RNA polymerase (RNAP) holoenzyme to activate transcription marks a milestone in deciphering the mechanism of transcriptional regulation by Wbls (6). σ^A^_4_ is highly conserved in the bacterial σ^70^-family primary sigma factors, responsible for recognizing the −35 promoter element and commonly used as an anchor for many transcription factors for gene activation (29–31). The subsequent studies show that all the *Mtb* Wbls, except for WhiB5, and two Wbl homologs in Streptomyces also bind to σ^A^_4_ (14,31–33), indicating a molecular mechanism shared by multiple Wbl members across different species.

Recent structural characterization of four Wbl proteins by our group and others reveals that, despite the low sequence identity, these Wbl proteins share a similar helical architecture surrounding the [4Fe-4S] binding pocket and bind to the same site on σ^A^_4_ centered on the H516-P517 pair (Fig. S1A and B) (26–28,34,35). Notably, the hydrophobic interactions-driven molecular interface within the Fe-S cluster binding pocket between these monomeric Wbls and σ^A^_4_ is distinct from other previously characterized σ^A^_4_-dependent transcription factors (*i.e*., the cyclic AMP receptor protein [CRP] and the fumarate and nitrate reduction regulatory protein [FNR]), which are dimeric and interact with σ^70^_4_ via surface charge complementarity (6,14,36). Further mechanistic investigations of *Mtb* WhiB7 and *Sve* WhiB2 show that WhiB7 activates gene expression by binding to σ^A^_4_ and specifically recognizing the target promoter sequence; while WhiB2 co-activates gene expression with another transcription factor, WhiA (26,27,34). However, a comprehensive analysis of the shared motifs and the mode of action in the Wbl family is currently missing.

By structural comparison of the reported σ^A^_4_-bound Wbl structures (26–28,32,34,35), including three Wbls (WhiB1, WhiB3 and WhiB7) from *Mtb* and the WhiB2 homolog from *Streptomyces venezuelae* (*Sve*), we found that Wbls binding to σ^A^_4_ is facilitated by a conserved patch of aromatic residues (herein referred to as the aromatic patch) in all four complexes, including one residue at the beginning of the helix *h1* (corresponding to W3 in *Mtb* WhiB1, labeled as <u>a</u>romatic <u>p</u>atch 1 [AP1]), two in the end of helix *h1* (corresponding to F17 and F18 in *Mtb* WhiB1 and labeled as AP2-3), and one in the β-turn connecting *h3* and *h4* (corresponding to W60 in *Mtb* WhiB1 and labeled as AP4) (Fig. 1A, Supplementary Fig. 1). The residues in the aromatic patch are conserved in each Wbl subfamily and at least three of them (including AP2-3) are critical for σ^A^_4_ binding and Wbl function (Fig. 1A, Supplementary Fig. 2) (26,28,35). In this study, we use multidisciplinary approaches to test whether the aromatic patch is a shared motif in the Wbl family for σ^A^_4_ binding. We present compelling structural, molecular and biochemical evidence demonstrating that the remaining *Mtb* Wbls (WhiB4, WhiB5 and WhiB6) also bind to the same site on σ^A^_4_, and the Wbl: σ^A^_4_ interaction is facilitated by the aromatic patch. Moreover, our structural modeling, phylogenetic analysis and biochemical characterization of Wbl proteins from diverse Actinobacteria and actinobacteriophages show that this aromatic patch is widely conserved among Wbl family members and uncover complex evolutionary relationships among bacterial and phage Wbl gene family members. Together, our findings identify a previously unrecognized structural feature that underlies a shared molecular mechanism of action in the Wbl family and offer new insights into the evolution and function of this unique family of [4Fe-4S] transcription factors.

## RESULTS

### Aromatic patch at the molecular interface of the WhiB6:σ^A^_4_ complex

WhiB6 is a Mycobacteria-specific Wbl protein. It regulates the expression of the Type VII secretion systems (ESX-1 and ESX-4) and plays an important role in the reactivation of *Mtb* and other pathogenic mycobacterial species (16,37–41). Compared to other *Mtb* Wbls, WhiB6 lacks two equivalent residues in the aromatic patch, and an unexpected polar residue is found corresponding to AP3, which has been shown to be essential for σ^A^_4_ binding in all the *Mtb* Wbls that we have tested thus far (Fig. 1A; Supplementary Fig. 2) (26,28,35). Due to the lack of any 3D structural information for either WhiB6 alone or in complex with σ^A^_4_, it is unclear how WhiB6 interacts with σ^A^_4_ for transcriptional regulation. To address this knowledge gap, we purified and crystallized the σ^A^_4_-bound WhiB6 to determine the crystal structure of the complex. As previously described, we used the chimera protein of σ^A^_4_ fused to the RNAP β-subunit flap tip helix (β_tip_) via an artificial linker (hereby denoted σ^A^_4_-β_tip_) to mimic the interaction between σ^A^ and the β-subunit in the RNAP holoenzyme (see in Materials and Methods, Supplementary Tables S1) (26). The crystal structure of WhiB6:σ^A^_4_-β_tip_ was refined at 1.8 Å (Supplementary Tables S2).

As observed in the other Wbl:σ^A^_4_ complexes (26–28,34,35), the overall architecture of WhiB6:σ^A^_4_ is dominated by alpha helices, and the molecular interface between σ^A^_4_ and WhiB6 is centered around the [4Fe-4S] cluster binding pocket (Fig. 1B). The core helical arrangement of WhiB6 surrounding the Fe-S cluster is surprisingly similar to that of other Wbl:σ^A^_4_ complexes despite the low sequence identity (<30%). A structural overlay of σ^A^_4_–bound WhiB6 with WhiB1 shows a Cα root mean square deviation (RMSD_Cα_) of 1.07 Å for the 37 aligned residues located in helices *h1-3* of WhiB1 (Fig. 1C). In contrast, striking differences are observed in the rest of the structural arrangements that are away from the Fe-S cluster binding pocket, in particular, WhiB6 has a significantly shorter helix *h2* and loop *l1* between *h1* and *h2*, but a much longer helix *h4* compared to WhiB1.

At the molecular interface of WhiB6:σ^A^_4_ complex, H516 and P517 of σ^A^_4_ interact with W21 and W56 of WhiB6 (corresponding to F17 [AP2] and W60 [AP4] of WhiB1) within the Fe-S cluster binding pocket of the complex (Fig. 1D). Additionally, a conserved Arg, R38, in the WhiB6 subfamily caps the Fe-S cluster pocket in the complex. R38 may serve as a functional equivalent of W3 in WhiB1 based on the structural comparison of the two complexes. As previously observed in other Wbl proteins, substitution of either H516 in σ^A^_4_ or W21 in WhiB6 by an Ala abolishes the complex formation in the pull-down assays (Fig. 1E and F), highlighting the essential role of the aromatic patch in the formation of WhiB6:σ^A^_4_. We note that the W21A mutation in WhiB6 disrupts the stability of the Fe-S cluster in WhiB6, as evidenced by a decrease in the absorption at ∼410 nm and the presence of the absorption peak at ∼350 nm when compared to the wildtype WhiB6. T22 of WhiB6 in loop *l1*, which corresponds to F18 of WhiB1 (Fig. 1D), is close to the molecular interface of the WhiB6:σ^A^_4_ complex within the [4Fe-4S] cluster binding pocket with the hydroxyl group of T22 facing away from the [4Fe-4S] cluster. Substitution of T22 with an Ala did not affect σ^A^_4_ binding in the pull-down assay, confirming a non-essential role of WhiB6 T22 for σ^A^_4_ binding and consistent with high sequence variability at this position among the WhiB6 homologs (Supplementary Fig. 1).

### WhiB5 binding to σ^A^_4_ is H516-dependent and facilitated by the aromatic patch

Unlike WhiB6, sequence analysis shows three of the aromatic patch residues are found in the WhiB5 subfamily, despite the low sequence identity (<30%) between WhiB5 and other *Mtb* Wbl proteins (Fig. 1A; Supplementary Fig. 2). Because of the observation, it was puzzling to note that WhiB5 is the only *Mtb* Wbl that did not show σ^A^ binding in the *in vitro* and *in vivo* binding assays by Feng *et al.* (31). However, as the authors pointed out, since the *in vitro* pull-down assays were carried out under aerobic conditions, oxidation damage of the [4Fe-4S] cluster in WhiB5 may also abolish σ^A^ binding. Therefore, we revisited the question of whether WhiB5 binds to σ^A^_4_. The results from our co-expression from *E. coli* and pull-down assays under anaerobic conditions confirm that WhiB5 binds to σ^A^_4_ and the binding requires σ^A^_4_ H516 (Fig. 2A), supporting that WhiB5 binds to the same site on σ^A^_4_ as other *Mtb* Wbls.

**Figure 2.**
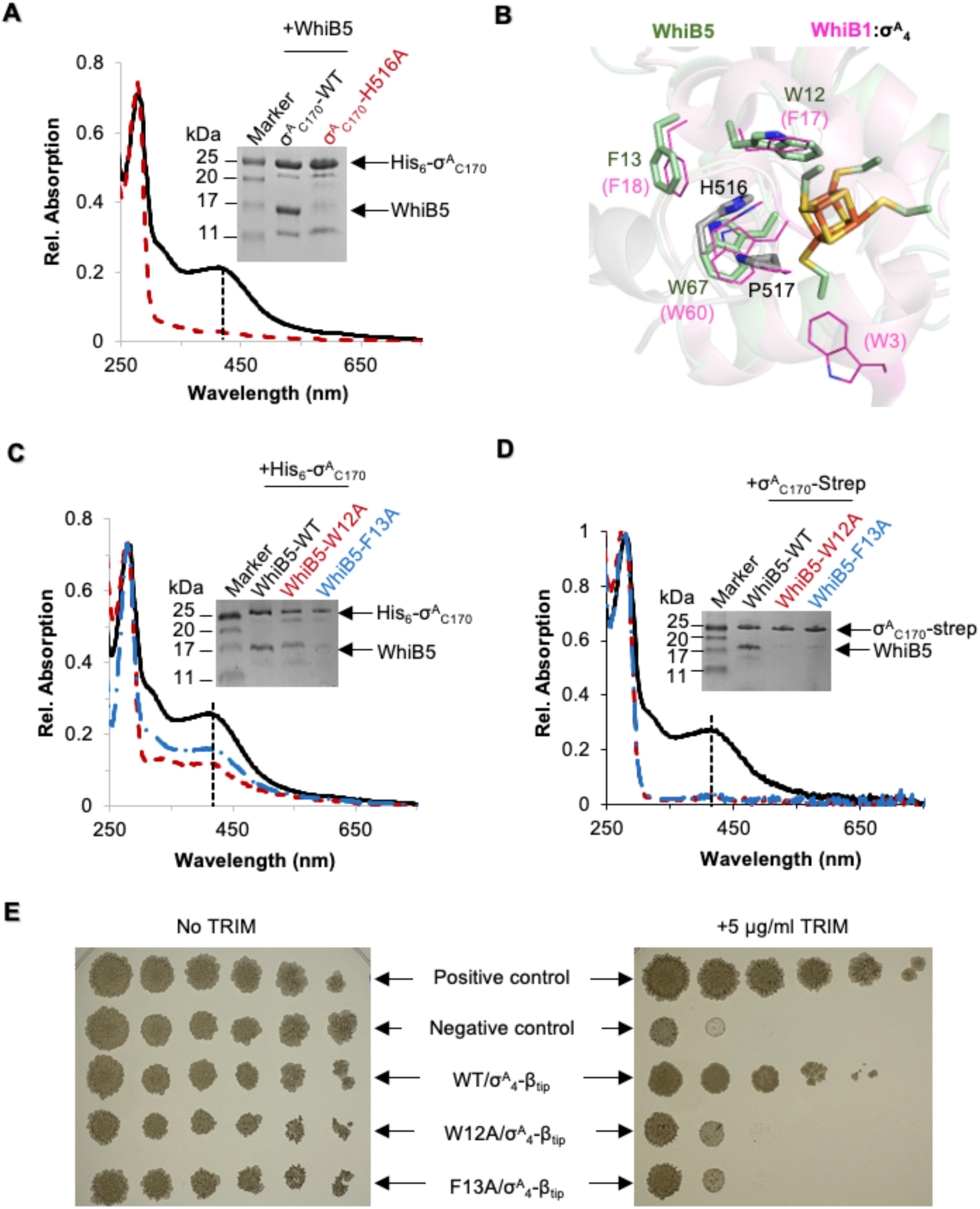
Characterization of WhiB5 binding to σ^A^_4_. (A) (C) and (D) UV-visible spectra and SDS-PAGE analyses of the samples from co-expression and affinity purification to test WhiB5 binding to σ^A^_4_ (wildtype or mutant as indicated). For Panels (A) and (C), His_6_-σ^A^_C170_ (wildtype or mutant as indicated) was used as the bait in the pull-down assays. For Panel (D), the Strep-tagged σ^A^_C170_ (σ^A^_C170_-Strep) was used as the bait. In all cases, the absorption spectra were normalized at 280 nm. The colors of the UV-Visible spectra correspond to those of the SDS-PAGE sample labels. The intensity of the absorption peak around 410 nm, highlighted with dashed lines, is indicative of the occupancy of the [4Fe-4S] cluster in the purified samples. The uncropped SDS PAGE images are shown in Supplementary Fig. 5. (B) Homology model of the σ^A^_4_-bound WhiB5 by SWISS-Model (pale green) overlaid with that of WhiB1 (PDB ID:6ONO, pink). The [4Fe-4S] clusters and the aromatic patch residues in Wbls, and the H516-P517 pair in σ^A^_4_ are shown in sticks. (E) Detection of the interactions between *Mtb* WhiB5 (wildtype and mutants as indicated) and σ^A^_4_ in *Mycobacterium smegmatis* (*Msm*) by Mycobacterial Protein Fragment Complementation (M-PFC) assay (see in Materials and Methods). Positive control, the *Msm* strain co-expressing the *Saccharomyces cerevisiae* leucine-zipper sequence homodimerization domain GCN4, with the fragment of mDHFR_F[1,2]_ and mDHFR_F[3]_, respectively, fused to the C-terminus of GCN4 (GCN4-mDHFR_F[1,2]_ and GCN4-mDHFR_F[3]_); Negative control, the *Msm* strain expressing the two fragments mDHFR_F[1,2]_ and mDHFR_F[3]_ separately. The *Msm* strain co-expressing the recombinant proteins, WhiB5-mDHFR_F[1,2]_ and σ^A^_C82_-β_tip_-mDHFR_F[3]_, to test the interaction between WhiB5 (wildtype or mutant as indicated) and σ^A^_4_. All the *Msm* strains were grown on the 7H10 broth agar plates in the absence or presence of 5 µg/ml trimethoprim (TRIM) as indicated. Cell growth in the presence of TRIM is indicative of the protein-protein interaction.

To gain structural insights into the interaction between WhiB5 and σ^A^_4_ and into the role of the aromatic patch residues in the interaction, we first attempted crystallographic characterization of WhiB5:σ^A^_4_ but were unable to produce crystals with this protein complex despite exhaustive trials. We therefore applied structural modeling approaches complemented by experimental validation to test the effects of the aromatic patch residues on WhiB5 binding to σ^A^_4_. both the homologous model of WhiB5:σ^A^_4_ by SWISS-MODEL and the *ab initio* model by AlphaFold3 support that WhiB5 adopts a similar helical arrangement as the σ^A^_4_-bound WhiB1 surrounding the cluster binding pocket (see in the Materials and Methods, Fig. 2A and Supplementary Fig. 4). The conserved aromatic patch residues in the WhiB5 subfamily align well with those in the WhiB1 at the molecular interface within the Fe-S binding pocket, implying a similar role in σ^A^_4_ binding. Consistent with the structural prediction, substitution of either W12 or F13 (corresponding to AP2 and AP3, respectively) by Ala significantly affects the cluster stability in WhiB5 and its binding to σ^A^_4_ in comparison to the wildtype WhiB5 in the pull-down assays (Fig. 2C-D). We note that a W12A or F13A mutation disrupts the stability of the Fe-S cluster in WhiB5, and we were not able to purify the mutant proteins in the holo-form. Low levels of WhiB5 mutants were detected by SDS PAGE analysis from the affinity pull-down samples using a His_6_-tagged σ^A^_4_ as the bait (Fig. 2C) but not in the samples using a Strep-tagged σ^A^_4_ as the bait (Fig. 2D), suggesting that the residual WhiB5 mutants retained in the pull-down samples from the NiNTA column are likely due to non-specific binding.

To assess whether WhiB5 binding to σ^A^_4_ is physiologically relevant, we examined the interaction between WhiB5 and σ^A^_4_ using the mycobacterial two-hybrid system assay termed mycobacterial protein fragment complementation (M-PFC) assay. The M-PFC assay is based on the functional reconstitution of two murine dihydrofolate reductase (mDHFR) domains, mDHFR_F[1,2]_ and mDHFR_F[3]_, respectively, by fusing the two domains independently to two interacting proteins. When the proteins of interest interact, mDHFR activity is restored, conferring bacterial resistance to trimethoprim (TRIM), an antibiotic targeting bacterial DHFR (42). The M-PFC assay was also used in the study by Feng *et al.* (31), in which mDHFR_F[1,2]_ was fused to the N-terminus of WhiB5. It is noted that WhiB5 has an unusually short sequence in the N-terminus prior to the first Fe-S cluster-ligating cysteine residue compared to other *Mtb* Wbl proteins (Fig. 1A). Indeed, we found that an N-terminal His-tag, but not a C-terminal tag, results in significant formation of inclusion bodies when over-expressing the recombinant WhiB5 protein in *E. coli* (data not shown). To avoid possible interference of WhiB5 folding and to stabilize the conformation of σ^A^_4_, we constructed the two mDHFR fragments to the C-terminus of WhiB5 and σ^A^_4_-β_tip_ (denoted WhiB5-mDHFR_F[1,2]_ and σ^A^_4_-mDHFR_F[3]_, respectively) in our M-PFC assay (see in Materials and Methods). As shown in Fig. 2E, all the strains grow similarly on the 7H10 agar plates without TRIM. In the presence of TRIM, the *Mycobacterium smegmatis* (*Msm*) strain expressing WhiB5-mDHFR_F[1,2]_ and σ^A^_4_-mDHFR_F[3]_, but not the negative control, grows on the TRIM agar plate comparably to the positive control. Consistent with the results from the pull-down assays (Fig. 2D), the *Msm* strain co-expressing σ^A^_4_-mDHFR_F[3]_ with either WhiB5 mutant (W12A or F13A)-mDHFR_F[1,2]_ shows significant defects in cell growth on the TRIM agar plate when compared to the wildtype WhiB5. These observations cement the physiological relevance of WhiB5 binding to σ^A^_4_ and underscore the essential role of the aromatic patch in WhiB5 for σ^A^_4_ binding.

Because both the *in vitro* and *in vivo* protein-protein binding assays were carried out with overexpressed proteins, we used isothermal titration calorimetry (ITC) to quantify WhiB5 binding to σ^A^_4_ under anaerobic conditions and compared the results with those of WhiB1 and WhiB2. WhiB1 and WhiB2 share the same aromatic patch compositions, making them good references for assessing the impact of the missing AP1 residue in WhiB5 on its binding affinity to σ^A^_4_. As shown in Fig. 3, all three Wbl proteins bind to σ^A^_4_ with low-nanomolar dissociation constants (*K_d_*) and the binding reactions are exothermic, which agrees favorably with the aromatic residues-driven interactions at the molecular interface of the Wbl:σ^A^_4_ complex and provides additional evidence supporting WhiB5 binding to σ^A^_4_ under physiologically relevant conditions. We note that the binding affinity of WhiB5 to σ^A^_4_ is about two- to three-fold lower than that of WhiB1 or WhiB2, consistent with the absence of a key aromatic patch residue (AP1) in WhiB5 for σ^A^_4_ binding (Fig. 1A).

**Figure 3.**
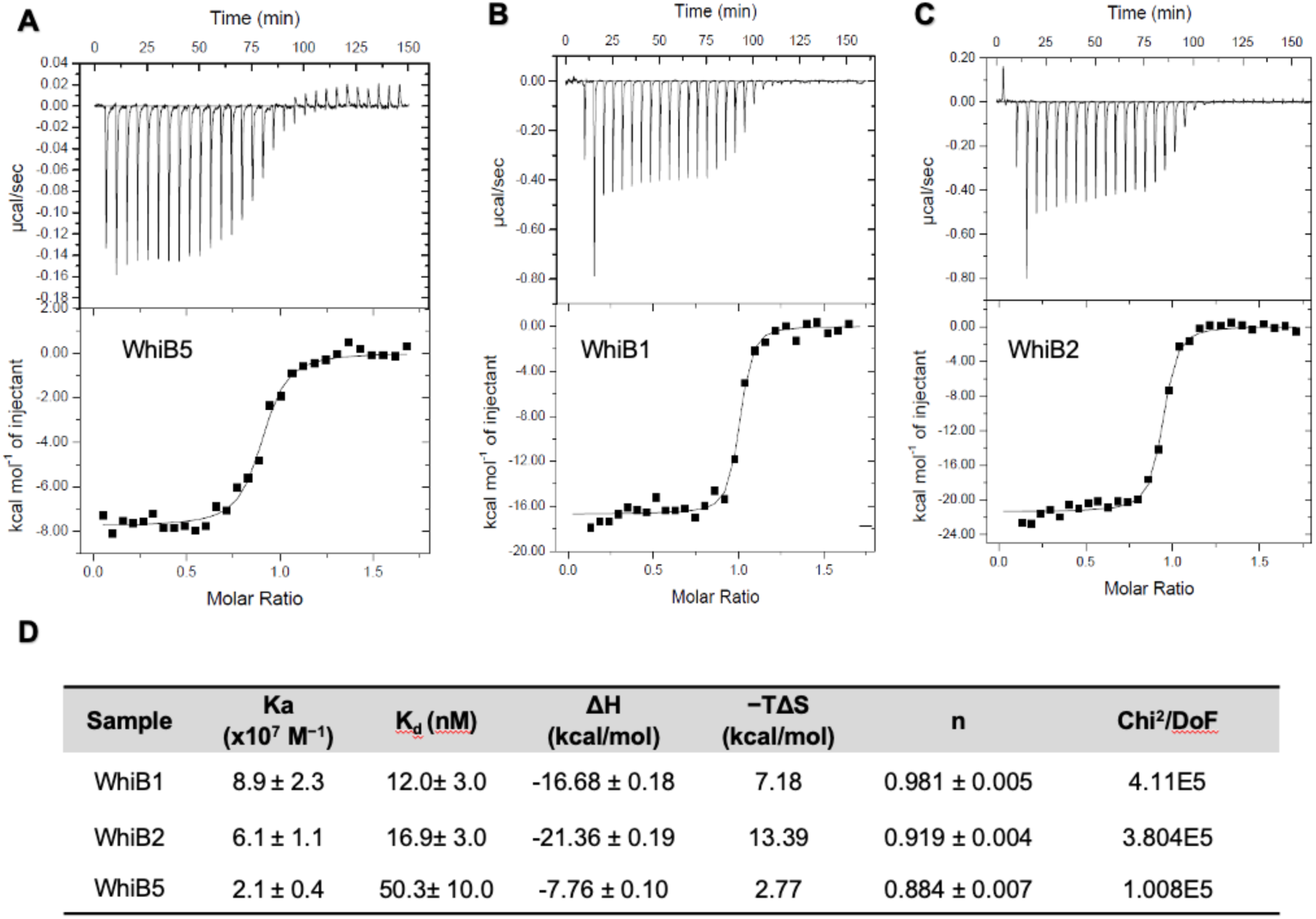
Isothermal titration calorimetry (ITC) assay of the interactions between σ^A^_4_ and *Mtb* Wbls. (A-C) Representative results for Isothermal titration calorimetry (ITC) assay of the interactions between σ^A^_4_ and *Mtb* Wbls as indicated. The calorimetric titration is shown in the top panel, with the integrated injection heat derived from the titration and the best-fit curve (dash lines) of a simple 1:1 interaction model shown in the bottom panel. (D) Summary of the binding parameters from the best fit, including the association constant (*K_a_*), dissociation constant (*K_d_*), changes in enthalpy (ΔH) and entropy (ΔS) upon binding, and the goodness of the fit (Chi^2^/DoF).

### Aromatic patch is required for WhiB4 binding to σ^A^_4_

Previous *in vitro* and *in vivo* protein–protein binding assays have shown that WhiB4 binds to σ^A^_4_, but the contribution of the aromatic patch residues to σ^A^_4_ binding has not been examined. Both the homology model and the AlphaFold model of WhiB4:σ^A^_4_ in our analysis predict that WhiB4 adapts a fold surrounding the Fe-S cluster resembling that of WhiB1, with the interaction interface centered on the H516-P517 pair of σ^A^_4_ (Supplementary Fig. 6). The aromatic patch residues in WhiB4 closely align with those of WhiB1, except for a leucine residue (L45) corresponding to AP2 (F17 in WhiB1) in the 3D structural alignment (Supplementary Fig. 6A).

Consistent with the structural prediction, our pull-down assays indicate that WhiB4 binding to σ^A^_4_ requires H516, as substitution of H516 by an Ala in σ^A^_4_ abolishes WhiB4 binding (Fig. 4A). Similarly, mutation of F46 (corresponding to F18 in WhiB1) to Ala in WhiB4 abolishes σ^A^_4_ binding and drastically destabilizes the WhiB4 [4Fe-4S] cluster, evident by a sharp decrease in the 410-nm absorbance (Fig. 4B). These observations underline the critical roles the aromatic patch residues in both maintaining [4Fe-4S] cluster integrity and binding to σ^A^_4_, as observed for other Wbls (Fig. 1F) (26,35). Notably, an Ala substitution of non-aromatic residue L45 also abolishes WhiB4 binding to σ^A^_4_. This observation differs from that of T22 in WhiB6 (Fig. 1E), but aligns well with the sequence analysis showing L45 is highly conserved in the WhiB4 subfamily and the structural models predicting that the side chain of L45 contributes directly to the WhiB4:σ^A^_4_ interface (Supplementary Figs. 2 and 6A).

**Figure 4.**
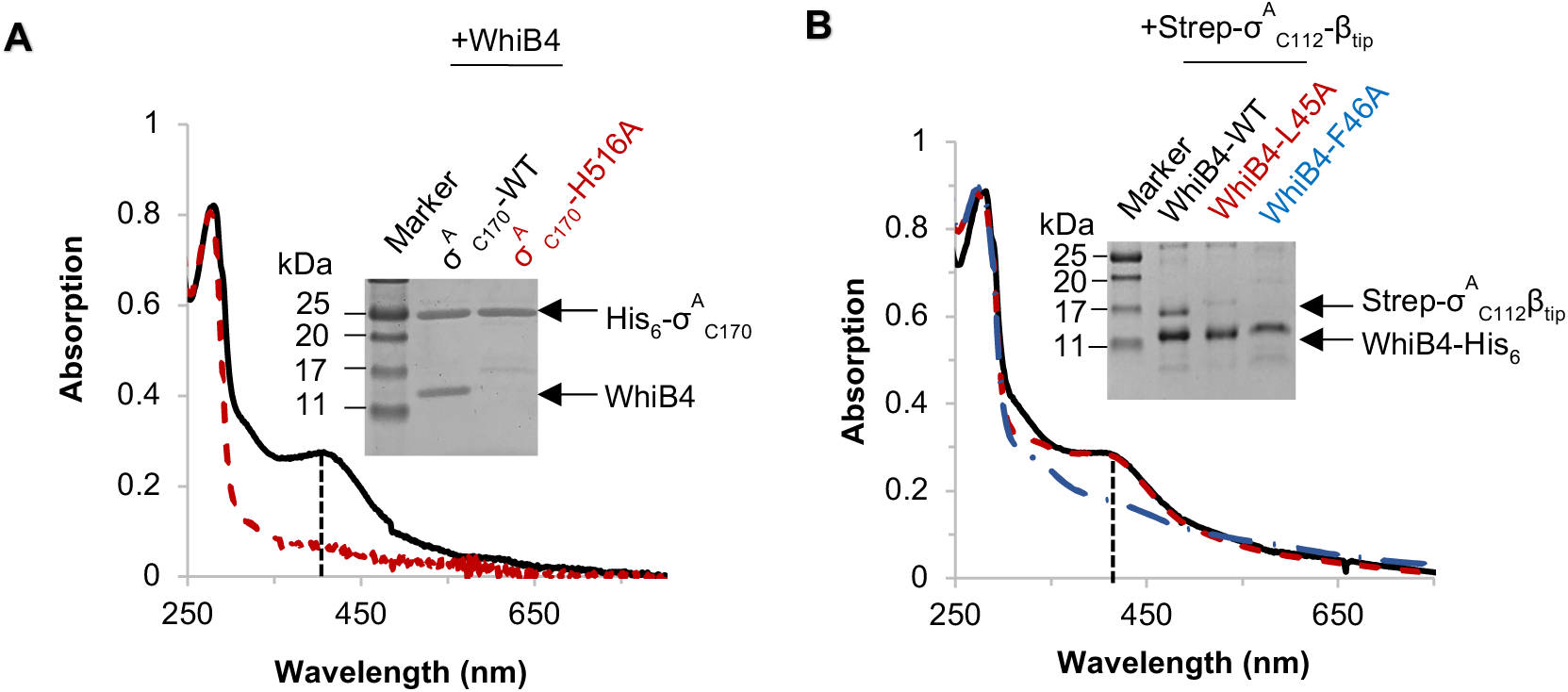
Role of the aromatic patch residues in WhiB4 binding to σ^A^_4._ (A) and (B) UV-visible spectra and SDS-PAGE analyses of the samples from co-expression and purification to test the interaction between σ^A^_4_ and WhiB4 (either wildtype or mutant as indicated). His_6_-σ^A^_C170_ (wildtype or mutant as indicated) was used as the bait in (A) to test the interaction with the tagless WhiB4. For Panel (B), WhiB4-His_6_ (wildtype or mutant as indicated) was used as the bait to test the interaction with Strep-σ^A^_4_-β_tip_. In both cases, the absorption spectra were normalized at 280 nm. The intensity of the absorption peak around 410 nm, highlighted with dashed lines, is indicative of the occupancy of the [4Fe-4S] cluster in the purified samples. The colors of the UV-Vis spectra match those of the SDS-PAGE sample labels. The uncropped SDS-PAGE images are shown in Supplementary Fig. 7.

### Aromatic patch facilitating Mtb Wbl binding to σ^A^_4_ is widely conserved in the Wbl family

To date, over 35,000 Wbl sequences have been deposited in the Wbl family database (Pfam ID: PF02467; InterPro ID: IPR034768). Since five *Mtb* Wbl proteins are well distributed in Actinobacteria and σ^A^_4_ is highly conserved in Actinobacteria, we speculate that the aromatic patch facilitating *Mtb* Wbls’ binding to σ^A^_4_ may be found in many other Wbl family proteins. To test this hypothesis, we analyzed 995 Wbl sequences from the representative proteomes of Actinobacteria and actinobacteriophages with a co-membership threshold of 15% (RP15), as defined in the Pfam database (Pfam ID: PF02467) (see in Materials and Methods). These Wbl proteins, of which 55% are from Actinobacteria and 45% are from Actinobacteriophages, were classified into 29 subfamilies (Fig. 5, Supplementary DataSet 1).

**Figure 5.**
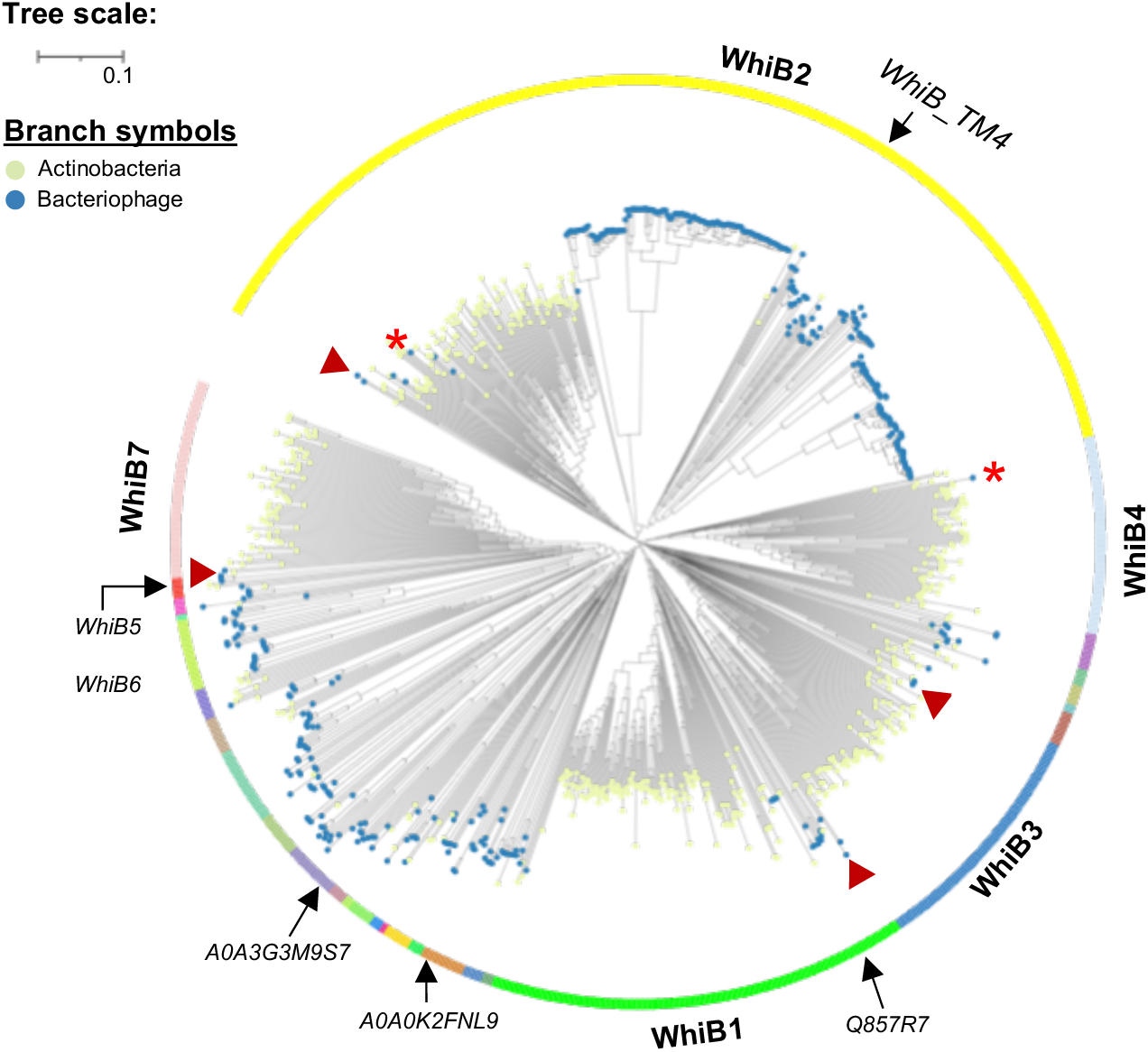
Phylogenetic analysis of the Wbl proteins from the representative genomes of Actinobacteria and actinobacteriophages. The phylogenetic tree of the 995 Wbl family proteins from the representative genomes of Actinobacteria and actinobacteriophages using the Pfam PF02467 (RP15), with highlights of the 29 Wbl subfamilies and the Wbls from actinobacteriophages. The five major Wbl subfamilies are labeled with the representative Wbls in *Mtb* (WhiB1-4 and WhiB7, respectively). The remaining smaller subfamilies are color-coded and assigned sequential numbers in the tree. Examples of the phage Wbl at the early diverging point of the clades are highlighted by the red arrowheads, while those nested within otherwise bacterial clades are highlighted by the red stars. The three phage Wbls (Uniprot IDs: A0A0K2FNL9, Q857R7 and A0A3G3M9S7) tested for σ^A^_4_ in this study are indicated on the tree.

Our phylogenetic analysis confirms that five *Mtb* Wbls (WhiB1–4 and WhiB7) represent the major Wbl subfamilies and reveals an intricate evolutionary relationship between bacterial Wbls and those from actinobacteriophages. As shown in Fig. 5, the five major Wbl subfamilies, marked by the representative *Mtb* Wbls, account for ∼80% of the total sequences from diverse actinobacterial species and associated phages, which is in line with the previous bioinformatic analysis (2). Among them, the WhiB2 subfamily is the largest Wbl subfamily and is found most widely distributed in Actinobacteria (2). Our analysis also shows that the WhiB2 subfamily has a broad mix of members from Actinobacteria (33%) and actinobacteriophages (67%), which include the WhiB2 homolog from Mycobacteriophage TM4 that has been shown to inhibit the function of WhiB2 in *Mycobacterium smegmatis* (*Msm*) (43). The other four major subfamilies are dominated by members from Actinobacteria. The relatively fewer actinobacteriophage sequences in these subfamilies are often located within the earliest diverging clades (highlighted by red arrowheads in Fig. 5), while the bacterial clades occupy more derived positions. This pattern suggests that many of these homologs were transferred from phages to bacteria, supporting the notion that the phages may have served as vectors for the horizontal gene transfer (HGT) of Wbl proteins among Actinobacteria as proposed in the previous studies (2,25,44). Conversely, the phylogenetic analysis also identifies some phage Wbl members nested within otherwise bacterial clades (red stars in Fig. 5), providing clear examples of HGT from bacteria to phage and underscoring the complex interplay of HGT in spreading *wbl* genes between Actinobacteria and their related phages. The 24 remaining subfamilies are much smaller in size, and the majority (19 out of 24) of them are dominated by Wbls from actinobacteriophages. The two remaining *Mtb* Wbls (WhiB5 and WhiB6) are grouped together with several unique Wbls from actinobacteriophages into one subfamily (labeled as No. 29 in Supplementary DataSet 1, Fig. 5), which is otherwise closely related with the WhiB7 subfamily.

As we speculated, sequence alignment of the 995 representative Wbl sequences indicates the aromatic patch residues in *Mtb* Wbls for σ^A^_4_ binding is well conserved among the Wbl homologs, together with the two known motifs (the four [4Fe-4S] cluster-ligating Cys and the β-turn) (Fig. 6A). Considering the low sequence identity among the Wbl members, we generated AlphaFold structural models for these Wbl members to better quantify the conservation of the aromatic patch in the Wbl family by 3D structural alignment against σ^A^_4_-bound WhiB1 (PDB ID:6ONO) (see in Materials and Methods). Our results show that, among the 995 sequences examined, 984 out of the 995 Wbls (98.7%) possess at least two aromatic residues (either Trp, Phe, Tyr or His) corresponding to AP1-4 and at least one corresponding to AP2-3 (Fig. 6B, Supplementary DataSet 1). Moreover, eight out of the 11 outliers possess an aromatic patch similar to that observed in the *Mtb* Wbl proteins when considering the residues next to the aromatic patch at the relative −1 to 1 position, with the remaining three lacking an aromatic patch as defined (Supplementary Fig. 8A-C). These findings indicate the newly identified aromatic patch is widely conserved in the Wbl family. During the structural comparison, we also found three Wbls with only three Cys at the expected [4Fe-4S] cluster binding pocket in the AlphaFold models. Two of these Wbl sequences (UniProt IDs: A0A1Q9SLC3 and A0A1Q9SWN8) are obsolete, while the third one (UniProt ID: A0A1D8EX64) from *Mycobacterium phage Tortellini* misses a Cys corresponding to the first Fe-S cluster-ligating Cys (corresponding to C9 in WhiB1) (Supplementary Fig. 8D). Blast search shows that A0A1D8EX64 shares 99% sequence identity with another Wbl protein (UniProt ID: A0A2U8UHK0) from *Mycobacterium phage Xavia*, which has an additional Cys in the N-terminus corresponding to C9 of WhiB1. Therefore, it is unclear whether these Wbl outliers are due to the uncertainty of gene sequencing or structural prediction, or they represent true evolutionary variants of the Wbl family.

**Figure 6.**
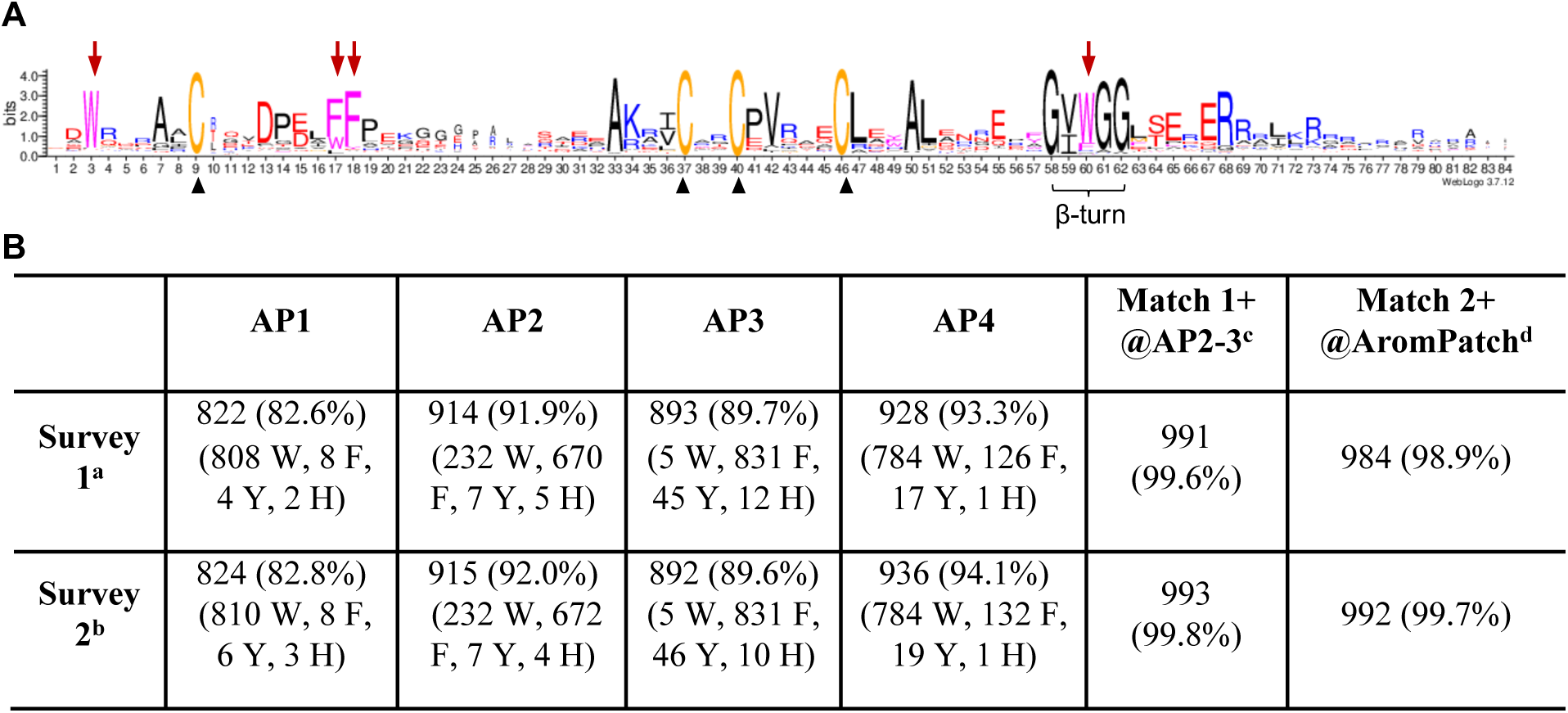
Conservation of the aromatic patch residues across the Wbl proteins from the representative genomes of Actinobacteria and actinobacteriophages. (A) The sequence logo of the 995 Wbl family proteins from the representative genomes of Actinobacteria and actinobacteriophages (Pfam PF02467, RP15) was generated by WebLogo. The four conserved cysteines for [4Fe-4S] cluster binding are highlighted by black arrowheads, and those in the aromatic patch indicated by red arrows. (B) Summary of the distribution of the aromatic patch in the Wbl family based on 3D structural alignment against *Mtb* WhiB1. ^a^Survey 1 includes the number of the aromatic residues (W, F, Y or H as indicated) corresponding to the aromatic patch residues AP1-4 (W3, F17, F18 and W60, respectively) in *Mtb* WhiB1 (PDB code:6ONO) by structural alignment. ^b^Survey 2 includes all the aromatic residues (W, F, Y or H as indicated) at the position −1 to +1 relative to the aromatic patch residues AP1-4 by structural alignment. ^c^Total number of Wbls containing at least one aromatic residue corresponding to AP2-3 (F17 or F18 in *Mtb* WhiB1). ^d^Total number of Wbls containing at least two aromatic residues corresponding to the aromatic patch in WhiB1.

To provide experimental evidence that Wbls from actinobacteriophages also bind to σ^A^_4_ in a manner similar to *Mtb* Wbls, we randomly selected three phage Wbls (UniProt IDs: A0A0K2FNL9, A0A3G3M9S7 and Q857R7) from the WhiB1 subfamily and two other relatively larger species-specific Wbl subfamilies (Fig. 5), and examined their interactions with σ^A^_4_. The pull-down assays confirm that, like the *Mtb* Wbls, all three phage Wbls bind to σ^A^_4_ in a His516-dependent manner (Supplementary Fig. 9). Moreover, as previously observed in *Mtb Wbls*, the aromatic patches in these phage Wbls vary in residue composition and differ in contribution to σ^A^_4_ binding and the [4Fe-4S] cluster stability. For example, Q857R7 belongs to the WhiB1 subfamily with an aromatic patch closely resembling that of WhiB1 (Fig. 5, Supplementary Fig. 10A). Consistently, the mutagenesis and pull-down assays show that the two aromatic patch residues F20 and F21 (corresponding to AP2 and AP3) are essential for σ^A^_4_ binding but do not affect the [4Fe-4S] cluster stability, mirroring the roles of F17 and F18 in WhiB1 (Supplementary Fig. 10B and C). In contrast, A0A3G3M9S7 shows low sequence identity (<25%) with *Mtb* WhiB1 and the AlphaFold model of σ^A^_4_-bound A0A3G3M9S7 does not align well with WhiB1 (RMSD_Cα_ of 2.68 Å for the 44 aligned residues; Supplementary Fig. 11A). Nonetheless, sequence alignment and homology modeling indicate that A0A3G3M9S7 retains three of the four WhiB1 aromatic patch residues (Supplementary Fig. 11B), with a lack of AP2 similar to WhiB4. Mutating either W12 or Y26 in A0A3G3M9S7 not only abolishes σ^A^_4_ binding but also destabilizes the [4Fe-4S] cluster as well as overall protein folding (Supplementary Fig. 11C-F). A0A0K2FNL9, on the other hand, contains three aromatic patch residues (W3, F17 or F18) aligning with AP1-3 in WhiB1 as well as two additional A0A3G3M9S7-spcific aromatic patch residues (W4 and F58) at the interaction interface of the A0A3G3M9S7:σ^A^_4_ model (Supplementary Fig. 12A). A single mutation of either AP2 or AP3 in A0A3G3M9S7 (F17A or F18A) or a double mutation of the A0A3G3M9S7-specific aromatic patch residues (W4A-F58A) has little effect on σ^A^_4_ binding in the pull-down assays (Supplementary Fig. 12B and C). However, a triple mutation (W4A-F17A-F58A) completely abolishes σ^A^_4_ binding and destabilizes the [4Fe-4S] cluster in A0A3G3M9S7. These observations parallel our previous finding that the WhiB3 subfamily-specific aromatic patch residue W15 compensates for the loss of the canonical aromatic patch residue AP1 (W17) in σ^A^_4_ binding (Supplementary Fig. 2) (28). Collectively, these results provide compelling experimental evidence that these phage Wbls bind to the same site on σ^A^_4_ facilitated by the aromatic patch and expand our knowledge of the diversity and adaptability of the aromatic patch in Wbl family proteins.

## Discussion

Wbl family proteins have gained increasing interest in the past decade because of their critical roles in the many biological processes in Actinobacteria that are of significant biomedical and economic implications, and their unique mode of action as the first family of monomeric Fe-S transcription factors. However, the mechanism of action of the Wbl family proteins remains enigmatic due to the lack of knowledge on the conserved functional motifs other than the [4Fe-4S] cluster binding motif and the β-turn. Herein, we apply multidisciplinary approaches to uncover a previously unrecognized motif, aromatic patch, in the Wbl family that facilitates σ^A^_4_ binding to regulate the genes involved in diverse biological processes and provide invaluable insights into the evolutionary history of the Wbl proteins among actinobacteria and associated phages. Multiple Wbl paralogs are present in *Mtb* and many other actinobacterial species, where they play nonredundant roles (4,22). Previous studies have shown that these paralogs are regulated at both the transcriptional and protein levels (8,12,45). Our earlier studies on WhiB1 and WhiB7 have demonstrated that the absence of an equivalent aromatic patch (AP) residue increases the oxygen sensitivity of the Wbl Fe–S cluster (26,35). In this study, we further show that the lack of an equivalent aromatic patch residue (AP1) in WhiB5 correlates with its lower binding affinity to σ^A^_4_. Together, these findings support an additional mechanism for modulating Wbl activities by fine-tuning the Fe-S reactivity and Wbl:σ^A^_4_ interaction.

The unusual hydrophobic residue-driven, tight interaction between Wbl proteins and σ^A^_4_ is in stark contrast to the previously characterized σ^70^_4_-dependent transcription activators that interact with σ^70^ via surface charge complementarity (6,14,36). Rather, its mode of σ^A^ binding resembles that of the “phage appropriator” AsiA of the bacteriophage T4, which hijacks the host RNAP via binding to σ^70^ and co-activates phage genes with another phage regulator MotA (46–48). Consistently, the phylogenetic analysis by us and other groups reveals a complex evolutionary relationship of Wbl homologs between actinobacteria and their associated phages. The *wbl* genes are believed to have been laterally distributed in Actinobacteria (44), while their origin remains unknown. Broad analyses of the Wbl family proteins indicate that some members are widespread among Actinobacteria, whereas others are more sporadically distributed (2). The variability in the distribution of multiple Wbl paralogs suggests that HGT may have played a role in their spread, and these transfer events were possibly mediated by phages, which also harbor Wbl genes (2,25,44). In fact, the Wbl family proteins were found to be among the most abundant regulator proteins encoded by actinobacteriophage genomes, and phylogenetic analysis revealed a complex evolutionary relationship of Wbl homologs between actinobacteria and their associated phages and plasmids, supporting a history of HGT (49), although the low sampling of phage homologs limits strong conclusions about the directionality of HGT. To further explore the evolutionary relationships among the bacterial and phage Wbl gene family members, our phylogenetic analysis included many additional phage Wbl sequences. Like previous studies, our tree shows a complex clustering of phage and bacterial Wbl homologs, lending further support for HGT of homologs between phage and bacteria. This finding implies that bacterial hosts may have sequestered the *wbl* gene from actinobacteriophages and repurposed the function of the genes for their own benefits during the early evolutionary events, followed by redistribution of the *wbl* genes among the Actinobacteria via phage-mediated HGT, in line with the previous hypothesis by Weinbauer *et al.* (50).

The results from this work and previous studies show that most Wbl characterized thus far, except for WhiB7, lack a recognizable DNA binding motif and do not bind to a specific target DNA sequence despite that they all bind to the same site on σ^A^_4_. These findings raise a new question about how Wbls regulate gene expression via binding to σ^A^ in the RNAP. Recent studies on the WhiB2 homolog from *Sve* offer a working model for how other Wbl proteins without a canonical DNA binding motif may regulate gene expression via binding to σ^A^ (15,34,51). These studies show that *Sve* WhiB co-activates the genes involved in cell division and sporulation via binding to σ^A^ in the RNAP and another regulator WhiA. The single-particle cryo-electron microscopy (cryoEM) structure of the WhiA/WhiB-dependent transcription initiation complex shows that *Sve* WhiB acts as a mediator that binds to WhiA on one side and σ^A^ on the other side (34). The WhiA and σ^A^_4_ in the RNAP, joined by WhiB2, define promoter specificity of the WhiA/WhiB2-dependent genes. *Sve* WhiB only shows non-specific DNA contacts, explaining the absence of the recognizable DNA binding motif in the WhiB2 subfamily. Further molecular and structural characterization of the Wbl-dependent transcriptional machinery is needed for an in-depth understanding of the diverse roles of Wbl proteins in stress-responsive transcriptional regulation.

## Methods

### Bacterial strains, plasmid, and growth conditions

The bacterial strains and plasmids used in this study are listed in supplementary Table S1. All the plasmids were confirmed by DNA sequencing before being transformed into *E. coli* BL21-Gold (DE3) strain for protein expression.

All *E. coli* strains were grown in Luria–Bertani (LB) media and at 37 °C, 200 rpm, unless otherwise specified. *Mycobacterium smegmatis* MC^2^ 155 (*Msm*) and the related strains were grown in either Middlebrook 7H9 broth or on 7H10 agar (BD Difco^TM^) supplemented with 10% v/v ADS (2% dextrose, 5% bovine serum albumin and 0.85% NaCl), 0.2% v/v glycerol and 0.05% v/v Tween 80 (Sigma). When appropriate, the media were supplemented with antibiotics at the following concentrations: ampicillin, 100 µg/ml; hygromycin, 100 µg/ml for *E. coli* and 50 µg/ml for *Msm*; kanamycin, 50 µg/ml for *E. coli* and 25 µg/ml for *Msm*; spectinomycin, 100 µg/ml for *E. coli*.

### General Procedures for protein overexpression and purification

The overexpression and purification of the proteins of interest in this study have been previously described unless otherwise stated (35). In brief, the *E. coli* BL21-Gold (DE3) strains containing the desirable plasmids were used to overexpress the proteins studied in this manuscript. Induction was initiated with the addition of 100 µM isopropyl β-d-thiogalactopyranoside (IPTG). For protein constructs containing a [4Fe-4S] cluster, 100 µM ferric ammonium sulfate was added before the induction.

The Wbl-containing proteins and complexes with a His-tag were purified using an affinity HisTrap HP column (GE Health Care Life Sciences) in an anaerobic chamber (Coy Labs). For proteins that tend to have high amounts of DNA contamination, an extra wash using a buffer containing 20 mM Tris-HCl, pH 8.0, 1M NaCl, and 1 mM DTT was added during the purification to reduce DNA contamination. Imidazole in the eluted fractions was removed by passing the eluate through either a desalting column or a Superdex 200 column (GE Healthcare Life Sciences). The recombinant σ^A^_4_-related proteins without a Wbl were purified with procedures similar to those for the purification of the Wbl proteins, except that the purification was carried out under aerobic conditions at 4°C.

The purified proteins were quantified by UV-visible (UV-Vis) spectroscopy and Bradford assays. The UV-Vis spectra of the purified proteins were recorded using an HP 8452a diode array UV-visible spectrophotometer (Agilent Technologies Inc.). The absorption at 410 nm, characteristic of proteins containing [4Fe-4S]^2+^ clusters (52,53), was used to estimate the occupancy of the Fe-S cluster in the protein samples containing a Wbl protein. Protein concentrations were estimated either by the Pierce Bradford Assay Kit (Thermo Fisher Scientific) or UV-visible absorption spectroscopy. The iron content in the purified proteins containing holo-Wbl was quantified as described (54). Unless otherwise specified, all the purified proteins in 50 mM Tris-HCl pH 8, 100 mM NaCl, and 1 mM DTT were stored in a Liquid Nitrogen Dewar or - 80°C until use.

### Crystallization

The plasmids pET21-MtbWhiB6 and pET28-His_6_-σ^A^_C82_β_tip_ were used to express and purify the WhiB6:σ^A^_C82_-β_tip_ complex. Initial crystallization screens of the WhiB6:σ^A^_C82_-β_tip_ complex were carried out at 18°C in a Coy anaerobic chamber using the sitting-drop vapor diffusion method, followed by optimization of the crystallization hits. High-quality crystals were obtained by mixing 1 μl of WhiB6:σ^A^_C82-_β_tip_ at 50 mg/ml in 50 mM Tris-HCl, pH 8.0, 100 mM NaCl and 1 mM DTT with an equal volume of the reservoir solution containing 0.1 M Na Citrate pH 5.5, 20% PEG3000. All the crystals were briefly soaked in the reservoir solution supplemented with 20% glycerol for cryoprotection before flash-frozen in liquid nitrogen.

### X-ray crystallographic data collection, structural determination, and analysis

X-ray diffraction data were collected at the beamline 12-2 of the Stanford Synchrotron Radiation Light source from single crystals maintained at 100 K using a 6M Pixel Array Detector. A single crystal was used to collect both the single anomalous dispersion (SAD) data at the Fe K-edge absorption peak (7140 eV), and the native data for final refinement was collected at 13,000 eV. The diffraction data were indexed, integrated, and scaled with HKL2000 (55). The experimental phases were determined by MR-SAD with Phenix.AutoSol using WhiB7:σ^A^_C82_-β_tip_ crystal structure (PDB: 7KUG) as a searching model (26,56). Model building and structure refinement were carried out using COOT and Phenix (57,58). Two copies of the WhiB6: σ^A^_C82_-β_tip_ complex molecules were found in each asymmetric unit. The data collection and refinement statistics are summarized in Table S2.

The gene sequence encoding 116-aa *Mtb* WhiB6 reported in the MycoBrowser portal was used to construct the plasmid for overexpression of the WhiB6:σ^A^_C82_-β_tip_ complex and for crystallization (59). We subsequently noticed that the two studies on the transcription analysis indicate the open reading frame of WhiB6 shifts 60 bp downstream, starting at M21 of WhiB6 reported in the MycoBrowser portal (60,61). We therefore renumbered the residue number accordingly.

### Co-expression and in vitro pull-down assays

To test the interaction between the Wbls protein (wildtype or mutant) and σ^A^_4_ (wildtype or mutant), the two plasmids encoding *Wbls* (wildtype or mutant as specified) and σ^A^_CTD_ (wildtype or mutant), respectively, were transformed into *E. coli* BL21-Gold (DE3) for protein co-expression and affinity purification as previously described (62). The plasmids used for the co-expression and pull-down assays are listed in Table S1. A HisTrap™ HP column was used for the pull-down assays using a His-tagged protein as the bait, while a Strep-Tactin^®^XT 4Flow^®^ column (IBA Lifesciences) was used for the pull-down assay using a Strep-tagged protein as the bait. The purified protein samples were analyzed by UV-Visible spectroscopy and SDS-PAGE.

### Isothermal titration calorimetry (ITC)

For isothermal titration calorimetry (ITC) experiments, the plasmids pET28-6Hisσ^A^_C112_-β_tip_, pETDuet-6His-SUMO-MtbWhiB1, pETDuet-6His-SUMO-MtbWhiB2 and pETDuet-6His-SUMO-MtbWhiB5 were used to express and purify 6His-tagged σ^A^_C112_-β_tip_ (His_6_-σ^A^_C112_-β_tip_) and the Wbl proteins with a N-terminal His_6_-SUMO tag, respectively, from *E. coli* as described above. Prior to the experiments, all the proteins were exchanged to the buffer containing 50 mM Tris pH 8.0, 100 mM NaCl, and 0.5 mM tris (2-carboxyethyl) phosphine (TCEP) by passing through a desalting column in an anaerobic chamber (Coy Labs).

The isothermal titration calorimetry (ITC) experiments were conducted at 25 °C in a VP-ITC calorimeter (MicroCal Malvern Panalytical LLC) inside an anaerobic chamber (Belle Technologies). For each experiment, 1.4 ml of 10 μM His_6_-σ^A^_C112_-β_tip_ was loaded into the sample cell. Following thermal equilibration, a series of 5-µl 75 μM His_6_-SUMO-Wbl was injected into the sample well at intervals of 300 seconds. A 2 μL pre-injection was included at the start of each titration. The reference buffer titrated to was used for background subtraction. The analysis for the ITC was done using MicroCal ITC Origin software.

### Mycobacterial Protein Fragment Complementation (M-PFC) assay

The interaction between WhiB5 and *Mtb* σ^A^_4_ was analyzed in *Msm* using the M-PFC assay (42). The plasmids used for the M-PFC assays are listed in Table S1. Briefly, the *Saccharomyces cerevisiae* leucine-zipper sequence homodimerization domain (GCN4), which was fused to the N terminus of dihydrofolate reductase fragment (mDHFR_F[1,2]_ and mDHFR_F[3]_) in pUAB100 and pUAB200 was replaced by WhiB5 or *Mtb* σ^A^_C82_-β_tip_, respectively, to express WhiB5-Gly_10_-mDHFR_F[1,2]_ and σ^A^_C82_-β_tip_-Gly_10_-mDHFR_F[3]_. The resulting plasmids, pUAB100-WhiB5 and pUAB200-σ^A^_C82_-β_tip_, were co-transformed into *Msm* for testing the interaction between WhiB5 and σ^A^_4_ in *Msm*. As previously described (42), the *Msm* strain carrying pUAB100 and pUAB200 was used as the positive control, while the strain with pUAB300 and pUAB400 was used as the negative control (42). All the strains were grown in 7H9 broth supplemented with 50 µg/ml of hygromycin and 25 µg/ml of kanamycin until saturation, followed by subculture with initial OD_600nm_ of 0.05, and incubated at 37 °C, 200 rpm until OD_600nm_ of 0.6-1. Each of the samples was diluted to OD_600nm_ of 0.4. 2 µl of the Msm cultures from ten-fold serial dilutions were spotted on 7H10 broth agar plates containing 50 µg/ml of hygromycin, 25 µg/ml of kanamycin, and trimethoprim (TRIM) as indicated. The images were taken after five days of incubation at 37 °C.

### Bioinformatic analysis

The 1048 Wbl sequences from the representative proteomes of Actinobacteria and actinobacteriophages with a co-membership threshold of 15% (RP15) were downloaded from the Pfam database (ID: PF02467) (63). We subsequently excluded the disqualified Wbl sequences, including those with less than four Cys residues per the definition of Wbl family or with non-standard amino acid (such as “X”). We also excluded the sequences outside of the range of 75–250 aa that are likely incomplete sequences or contain multiple domains. The resulting 995 sequences were used in the following bioinformatic analysis (Supplementary DataSet 1).

For the phylogenetic analysis, multiple sequence alignment of the resulting 995 sequences was conducted by ClustalW (64). Fasttree2 and TreeCluster were used for tree structure generation and cluster assignment, respectively (65,66). The classing method (avg clade) in the TreeCluster program was used for the cluster assignment. The five major clusters were named after the Mtb Wbl proteins in the cluster, while the automatically assigned cluster IDs remain for the rest of the clusters.

### 3D structural modeling and alignment

AlphaFold2 was used for the structural modeling of the representative Wbl proteins described above, with the template data up to 2022 (67). The top-ranked AlphaFold model for each Wbl sequence was used for 3D structural alignment against WhiB1 (PDB ID: 6ONO) by the PyMol Molecular Graphics System v3.0 (https://pymol.org).

The homologous models of the Wbl:σ^A^_4_ complexes were generated by SWISS-MODEL using WhiB1:σ^A^_4_ (PDB ID: 6ONO) as a template, while the *ab initio* models of the complexes were generated by AlphaFold3 using the online server (35,68,69).

### Data visualization

The representative homologs for the five major Wbl subfamilies (WhiB1-4 and WhiB7) used for the alignments are modified from the study by *Chandra, et al.* (2), and listed in the legend of Figure S1. The homologs of WhiB5 and WhiB6 are selected from the represented mycobacterial species as noted in the legend of Figure S1. Sequence alignments were performed using Clustal Omega and the ESpript online server (https://espript.ibcp.fr), and the sequence logo was generated using WebLogo (70–72).

The phylogenetic tree structure and protein clusters from the bioinformatic analysis were visualized by iTOL (73). 3D structure figures were prepared with the PyMol Molecular Graphics System v3.0 (https://pymol.org).

## Supporting information

Validation report

## Data Availability

Atomic coordinates and structure factors have been deposited in the RCSB Protein Data Bank (PDB) under the accession code 8D5V for the WhiB6:σ^A^_4_-β_tip_ complex, and released for public access with the validation report.

The [phylogenetic tree](https://itol.embl.de/tree/12993161230227991746751886) with highlights of the Wbl subfamilies and the actinobacteriophages Wbls is available at iTOL.

## Supplemental Information

Supplemental documents can be found online at the journal website.

## Acknowledgments

The authors thank Dr. Mark A. Wilson at the University of Nebraska-Lincoln for the insightful discussions. We thank the staff at beamlines 12-2 of SSRL for their assistance during the X-ray diffraction data collection. The authors would like to acknowledge the following funding sources: This work was supported by grants from the National Institutes of Health (R35 GM138157) and the National Science Foundation CAREER Award (CLP 1846908 for the work related to WhiB1) to L-M.Z., and the NIH (T32 GM136593) to D.G.B. through the T32 Predoctoral Training Fellowship. The content is solely the responsibility of the authors and does not necessarily represent the official views of the National Institutes of Health and the National Science Foundation. Use of the Stanford Synchrotron Radiation Lightsource, SLAC National Accelerator Laboratory, is supported by the U.S. Department of Energy, Office of Science, Office of Basic Energy Sciences under Contract No. DE-AC02-76SF00515. The SSRL Structural Molecular Biology Program is supported by the DOE Office of Biological and Environmental Research, and by the National Institutes of Health, National Institute of General Medical Sciences (including P41GM103393). This work was completed utilizing the Holland Computing Center of the University of Nebraska, which receives support from the UNL Office of Research and Innovation and the Nebraska Research Initiative.

## Author contributions

L-M.Z. directed the project. D.G.B., T.W., C. J., S. L., M.H and Z. L. performed molecular biology work, expressed, purified, and characterized the proteins. T.W. carried out the crystallographic study. J.S. and D.G.B. conducted the ITC assay, with assistance from D.B. Y.P., H.Y, D.G.B. and L-M.Z. carried out the structural modeling and structural analysis. Q.Y., P. M. and C.O. conducted the phylogenetic analysis and tree figure generation, J.P.M. provided expertise in the phylogenetic tree interpretation. A. S. provided the plasmids used for M-PFC assays. D.G.B, T.W, J.P.M. and L-M.Z. wrote the manuscript. All authors contributed to the manuscript preparation.

## References

1. Chater, K. F. (1972) A morphological and genetic mapping study of white colony mutants of *Streptomyces coelicolor*. J Gen Microbiol 72, 9–28

2. Chandra, G., and Chater, K. F. (2014) Developmental biology of Streptomyces from the perspective of 100 actinobacterial genome sequences. FEMS Microbiol Rev 38, 345–379

3. Davis, N. K., and Chater, K. F. (1992) The *Streptomyces coelicolor* whiB gene encodes a small transcription factor-like protein dispensable for growth but essential for sporulation. Mol Gen Genet 232, 351–358

4. Bush, M. J. (2018) The actinobacterial WhiB-like (Wbl) family of transcription factors. Mol Microbiol 110, 663–676

5. Molle, V., Palframan, W. J., Findlay, K. C., and Buttner, M. J. (2000) WhiD and WhiB, homologous proteins required for different stages of sporulation in *Streptomyces coelicolor A3(2)*. J Bacteriol 182, 1286–1295

6. Steyn, A. J., Collins, D. M., Hondalus, M. K., Jacobs, W. R., Jr., Kawakami, R. P., and Bloom, B. R. (2002) *Mycobacterium tuberculosis* WhiB3 interacts with RpoV to affect host survival but is dispensable for *in vivo* growth. Proc Natl Acad Sci USA 99, 3147–3152

7. Morris, R. P., Nguyen, L., Gatfield, J., Visconti, K., Nguyen, K., Schnappinger, D., Ehrt, S., Liu, Y., Heifets, L., Pieters, J., Schoolnik, G., and Thompson, C. J. (2005) Ancestral antibiotic resistance in *Mycobacterium tuberculosis*. Proc Natl Acad Sci USA 102, 12200–12205

8. Geiman, D. E., Raghunand, T. R., Agarwal, N., and Bishai, W. R. (2006) Differential gene expression in response to exposure to antimycobacterial agents and other stress conditions among seven *Mycobacterium tuberculosis* whiB-like genes. Antimicrob Agents Chemother 50, 2836–2841

9. Singh, A., Guidry, L., Narasimhulu, K. V., Mai, D., Trombley, J., Redding, K. E., Giles, G. I., Lancaster, J. R., Jr., and Steyn, A. J. (2007) *Mycobacterium tuberculosis* WhiB3 responds to O_2_ and nitric oxide via its [4Fe-4S] cluster and is essential for nutrient starvation survival. Proc Natl Acad Sci USA 104, 11562–11567

10. Smith, L. J., Stapleton, M. R., Fullstone, G. J., Crack, J. C., Thomson, A. J., Le Brun, N. E., Hunt, D. M., Harvey, E., Adinolfi, S., Buxton, R. S., and Green, J. (2010) *Mycobacterium tuberculosis* WhiB1 is an essential DNA-binding protein with a nitric oxide-sensitive iron-sulfur cluster. Biochem J 432, 417–427

11. Chawla, M., Parikh, P., Saxena, A., Munshi, M., Mehta, M., Mai, D., Srivastava, A. K., Narasimhulu, K. V., Redding, K. E., Vashi, N., Kumar, D., Steyn, A. J., and Singh, A. (2012) Mycobacterium tuberculosis WhiB4 regulates oxidative stress response to modulate survival and dissemination in vivo. Mol Microbiol 85, 1148–1165

12. Larsson, C., Luna, B., Ammerman, N. C., Maiga, M., Agarwal, N., and Bishai, W. R. (2012) Gene expression of *Mycobacterium tuberculosis* putative transcription factors *whiB1-7* in redox environments. PLoS One 7, e37516

13. Casonato, S., Cervantes Sanchez, A., Haruki, H., Rengifo Gonzalez, M., Provvedi, R., Dainese, E., Jaouen, T., Gola, S., Bini, E., Vicente, M., Johnsson, K., Ghisotti, D., Palu, G., Hernandez-Pando, R., and Manganelli, R. (2012) WhiB5, a transcriptional regulator that contributes to *Mycobacterium tuberculosis* virulence and reactivation. Infect Immun 80, 3132–3144

14. Burian, J., Yim, G., Hsing, M., Axerio-Cilies, P., Cherkasov, A., Spiegelman, G. B., and Thompson, C. J. (2013) The mycobacterial antibiotic resistance determinant WhiB7 acts as a transcriptional activator by binding the primary sigma factor SigA (RpoV). Nucleic Acids Res 41, 10062–10076

15. Bush, M. J., Chandra, G., Bibb, M. J., Findlay, K. C., and Buttner, M. J. (2016) Genome-wide chromatin immunoprecipitation sequencing analysis shows that WhiB is a transcription factor that cocontrols its regulon with WhiA to initiate developmental cell division in *Streptomyces*. mBio 7, e00523–00516

16. Chen, Z., Hu, Y., Cumming, B. M., Lu, P., Feng, L., Deng, J., Steyn, A. J., and Chen, S. (2016) Mycobacterial WhiB6 differentially regulates ESX-1 and the Dos regulon to modulate granuloma formation and virulence in Zebrafish. Cell Rep 16, 2512–2524

17. Bosserman, R. E., Nguyen, T. T., Sanchez, K. G., Chirakos, A. E., Ferrell, M. J., Thompson, C. R., Champion, M. M., Abramovitch, R. B., and Champion, P. A. (2017) WhiB6 regulation of ESX-1 gene expression is controlled by a negative feedback loop in *Mycobacterium marinum*. Proc Natl Acad Sci USA 114, E10772–E10781

18. Wu, J., Ru, H. W., Xiang, Z. H., Jiang, J., Wang, Y. C., Zhang, L., and Liu, J. (2017) WhiB4 Regulates the PE/PPE Gene Family and is Essential for Virulence of Mycobacterium marinum. Sci Rep 7, 3007

19. Lee, D. S., Kim, P., Kim, E. S., Kim, Y., and Lee, H. S. (2018) *Corynebacterium glutamicum* WhcD interacts with WhiA to exert a regulatory effect on cell division genes. Antonie Van Leeuwenhoek 111, 641–648

20. Chawla, M., Mishra, S., Anand, K., Parikh, P., Mehta, M., Vij, M., Verma, T., Singh, P., Jakkala, K., Verma, H. N., AjitKumar, P., Ganguli, M., Narain Seshasayee, A. S., and Singh, A. (2018) Redox-dependent condensation of the mycobacterial nucleoid by WhiB4. Redox Biol 19, 116–133

21. Lin, C., Tang, Y., Wang, Y., Zhang, J., Li, Y., Xu, S., Xia, B., Zhai, Q., Li, Y., Zhang, L., and Liu, J. (2022) WhiB4 Is Required for the Reactivation of Persistent Infection of *Mycobacterium marinum* in Zebrafish. Microbiol Spectr 10, e0044321

22. Guiza Beltran, D., Wan, T., and Zhang, L. (2024) WhiB-like proteins: Diversity of structure, function and mechanism. Biochim Biophys Acta Mol Cell Res 1871, 119787

23. Soliveri, J. A., Gomez, J., Bishai, W. R., and Chater, K. F. (2000) Multiple paralogous genes related to the Streptomyces coelicolor developmental regulatory gene *whiB* are present in Streptomyces and other actinomycetes. Microbiology 146 (Pt 2), 333–343

24. Burian, J., Ramon-Garcia, S., Sweet, G., Gomez-Velasco, A., Av-Gay, Y., and Thompson, C. J. (2012) The mycobacterial transcriptional regulator *whiB7* gene links redox homeostasis and intrinsic antibiotic resistance. J Biol Chem 287, 299–310

25. Saini, V., Farhana, A., and Steyn, A. J. (2012) *Mycobacterium tuberculosis* WhiB3: a novel iron-sulfur cluster protein that regulates redox homeostasis and virulence. Antioxid Redox Signal 16, 687–697

26. Wan, T., Horova, M., Beltran, D. G., Li, S., Wong, H. X., and Zhang, L. M. (2021) Structural insights into the functional divergence of WhiB-like proteins in *Mycobacterium tuberculosis*. Mol Cell 81, 2887–2900

27. Lilic, M., Darst, S. A., and Campbell, E. A. (2021) Structural basis of transcriptional activation by the *Mycobacterium tuberculosis* intrinsic antibiotic-resistance transcription factor WhiB7. Mol Cell 81, 2875–2886 e2875

28. Wan, T., Horova, M., Khetrapal, V., Li, S., Jones, C., Schacht, A., Sun, X., and Zhang, L. (2023) Structural basis of DNA binding by the WhiB-like transcription factor WhiB3 in *Mycobacterium tuberculosis*. The Journal of biological chemistry 299, 104777

29. Gomez, M., Doukhan, L., Nair, G., and Smith, I. (1998) sigA is an essential gene in Mycobacterium smegmatis. Molecular Microbiology 29, 617–628

30. Lee, D. J., Minchin, S. D., and Busby, S. J. (2012) Activating transcription in bacteria. Annu Rev Microbiol 66, 125–152

31. Feng, L., Chen, Z., Wang, Z., Hu, Y., and Chen, S. (2016) Genome-wide characterization of monomeric transcriptional regulators in *Mycobacterium tuberculosis*. Microbiology 162, 889–897

32. Kudhair, B. K., Hounslow, A. M., Rolfe, M. D., Crack, J. C., Hunt, D. M., Buxton, R. S., Smith, L. J., Le Brun, N. E., Williamson, M. P., and Green, J. (2017) Structure of a Wbl protein and implications for NO sensing by *M. tuberculosis*. Nat Commun 8, 2280

33. Stewart, M. Y. Y., Bush, M. J., Crack, J. C., Buttner, M. J., and Le Brun, N. E. (2020) Interaction of the Streptomyces Wbl protein WhiD with the principal sigma factor σ^HrdB^ depends on the WhiD [4Fe-4S] cluster. J Biol Chem 295, 9752–9765

34. Lilic, M., Holmes, N. A., Bush, M. J., Marti, A. K., Widdick, D. A., Findlay, K. C., Choi, Y. J., Froom, R., Koh, S., Buttner, M. J., and Campbell, E. A. (2023) Structural basis of dual activation of cell division by the actinobacterial transcription factors WhiA and WhiB. Proc Natl Acad Sci USA 120, e2220785120

35. Wan, T., Li, S., Beltran, D. G., Schacht, A., Zhang, L., Becker, D. F., and Zhang, L. (2020) Structural basis of non-canonical transcriptional regulation by the σ^A^-bound iron-sulfur protein WhiB1 in *M. tuberculosis*. Nucleic acids research 48, 501–516

36. Lonetto, M. A., Rhodius, V., Lamberg, K., Kiley, P., Busby, S., and Gross, C. (1998) Identification of a contact site for different transcription activators in region 4 of the *Escherichia coli* RNA polymerase σ^70^ subunit. J Mol Biol 284, 1353–1365

37. Abdallah, A. M., Weerdenburg, E. M., Guan, Q., Ummels, R., Borggreve, S., Adroub, S. A., Malas, T. B., Naeem, R., Zhang, H., Otto, T. D., Bitter, W., and Pain, A. (2019) Integrated transcriptomic and proteomic analysis of pathogenic mycobacteria and their esx-1 mutants reveal secretion-dependent regulation of ESX-1 substrates and WhiB6 as a transcriptional regulator. PLoS One 14, e0211003

38. Bosserman, R. E., Nguyen, T. T., Sanchez, K. G., Chirakos, A. E., Ferrell, M. J., Thompson, C. R., Champion, M. M., Abramovitch, R. B., and Champion, P. A. (2017) WhiB6 regulation of ESX-1 gene expression is controlled by a negative feedback loop in Mycobacterium marinum. Proc Natl Acad Sci U S A 114, E10772–E10781

39. Solans, L., Aguilo, N., Samper, S., Pawlik, A., Frigui, W., Martin, C., Brosch, R., and Gonzalo-Asensio, J. (2014) A specific polymorphism in Mycobacterium tuberculosis H37Rv causes differential ESAT-6 expression and identifies WhiB6 as a novel ESX-1 component. Infect Immun 82, 3446–3456

40. Sala, C., Odermatt, N. T., Soler-Arnedo, P., Gulen, M. F., von Schultz, S., Benjak, A., and Cole, S. T. (2018) EspL is essential for virulence and stabilizes EspE, EspF and EspH levels in Mycobacterium tuberculosis. PLoS Pathog 14, e1007491

41. Vijayaraj, M., Abhinand, P. A., and Ragunath, P. K. (2019) Virtual screening of a MDR-TB WhiB6 target identified by gene expression profiling. Bioinformation 15, 557–567

42. Singh, A., Mai, D., Kumar, A., and Steyn, A. J. (2006) Dissecting virulence pathways of *Mycobacterium tuberculosis* through protein-protein association. Proc Natl Acad Sci USA 103, 11346–11351

43. Rybniker, J., Nowag, A., van Gumpel, E., Nissen, N., Robinson, N., Plum, G., and Hartmann, P. (2010) Insights into the function of the WhiB-like protein of mycobacteriophage TM4 – a transcriptional inhibitor of WhiB2. Mol Microbiol 77, 642–657

44. Chater, K. F., and Chandra, G. (2006) The evolution of development in Streptomyces analysed by genome comparisons. FEMS Microbiol Rev 30, 651–672

45. Raju, R. M., Jedrychowski, M. P., Wei, J. R., Pinkham, J. T., Park, A. S., O’Brien, K., Rehren, G., Schnappinger, D., Gygi, S. P., and Rubin, E. J. (2014) Post-translational regulation via Clp protease is critical for survival of *Mycobacterium tuberculosis*. PLoS Pathog 10, e1003994

46. Lambert, L. J., Wei, Y., Schirf, V., Demeler, B., and Werner, M. H. (2004) T4 AsiA blocks DNA recognition by remodeling σ^70^ region 4. EMBO J 23, 2952–2962

47. Gregory, B. D., Nickels, B. E., Garrity, S. J., Severinova, E., Minakhin, L., Urbauer, R. J. B., Urbauer, J. L., Heyduk, T., Severinov, K., and Hochschild, A. (2004) A regulator that inhibits transcription by targeting an intersubunit interaction of the RNA polymerase holoenzyme. Proc Natl Acad Sci USA 101, 4554–4559

48. Shi, J., Wen, A., Zhao, M., You, L., Zhang, Y., and Feng, Y. (2019) Structural basis of sigma appropriation. Nucleic Acids Res 47, 9423–9432

49. Sharma, V., Hardy, A., Luthe, T., and Frunzke, J. (2021) Phylogenetic Distribution of WhiB- and Lsr2-Type Regulators in Actinobacteriophage Genomes. Microbiol Spectr 9, e0072721

50. Weinbauer, M. G., and Rassoulzadegan, F. (2004) Are viruses driving microbial diversification and diversity? Environ Microbiol 6, 1–11

51. Bush, M. J., Bibb, M. J., Chandra, G., Findlay, K. C., and Buttner, M. J. (2013) Genes Required for Aerial Growth, Cell Division, and Chromosome Segregation Are Targets of WhiA before Sporulation in *Streptomyces venezuelae*. mBio 4

52. Johnson, M. K. (1998) Iron-sulfur proteins: new roles for old clusters. Curr Opin Chem Biol 2, 173–181

53. Crack, J. C., Smith, L. J., Stapleton, M. R., Peck, J., Watmough, N. J., Buttner, M. J., Buxton, R. S., Green, J., Oganesyan, V. S., Thomson, A. J., and Le Brun, N. E. (2011) Mechanistic insight into the nitrosylation of the [4Fe-4S] cluster of WhiB-like proteins. J Am Chem Soc 133, 1112–1121

54. Crack, J. C., Green, J., Thomson, A. J., and Le Brun, N. E. (2014) Techniques for the production, isolation, and analysis of iron-sulfur proteins. Methods Mol Biol 1122, 33–48

55. Otwinowski, Z., and Minor, W. (1997) Processing of X-ray diffraction data collected in oscillation mode. Methods Enzymol 276, 307–326

56. Adams, P. D., Afonine, P. V., Bunkoczi, G., Chen, V. B., Davis, I. W., Echols, N., Headd, J. J., Hung, L. W., Kapral, G. J., Grosse-Kunstleve, R. W., McCoy, A. J., Moriarty, N. W., Oeffner, R., Read, R. J., Richardson, D. C., Richardson, J. S., Terwilliger, T. C., and Zwart, P. H. (2010) PHENIX: a comprehensive Python-based system for macromolecular structure solution. Acta Crystallogr D 66, 213–221

57. Adams, P. D., Afonine, P. V., Bunkoczi, G., Chen, V. B., Davis, I. W., Echols, N., Headd, J. J., Hung, L. W., Kapral, G. J., Grosse-Kunstleve, R. W., McCoy, A. J., Moriarty, N. W., Oeffner, R., Read, R. J., Richardson, D. C., Richardson, J. S., Terwilliger, T. C., and Zwart, P. H. (2010) PHENIX: a comprehensive Python-based system for macromolecular structure solution. Acta Crystallogr D Biol Crystallogr 66, 213–221

58. Emsley, P., Lohkamp, B., Scott, W. G., and Cowtan, K. (2010) Features and development of Coot. Acta Crystallogr D Biol Crystallogr 66, 486–501

59. Kapopoulou, A., Lew, J. M., and Cole, S. T. (2011) The MycoBrowser portal: a comprehensive and manually annotated resource for mycobacterial genomes. Tuberculosis (Edinb) 91, 8–13

60. Shell, S. S., Wang, J., Lapierre, P., Mir, M., Chase, M. R., Pyle, M. M., Gawande, R., Ahmad, R., Sarracino, D. A., Ioerger, T. R., Fortune, S. M., Derbyshire, K. M., Wade, J. T., and Gray, T. A. (2015) Leaderless Transcripts and Small Proteins Are Common Features of the Mycobacterial Translational Landscape. PLoS Genet 11, e1005641

61. Cortes, T., Schubert, Olga T., Rose, G., Arnvig, Kristine B., Comas, I., Aebersold, R., and Young, Douglas B. (2013) Genome-wide mapping of transcriptional start sites defines an extensive leaderless transcriptome in *Mycobacterium tuberculosis*. Cell Reports 5, 1121–1131

62. Wan, T., Horova, M., Beltran, D. G., Li, S., Wong, H. X., and Zhang, L. M. (2021) Structural insights into the functional divergence of WhiB-like proteins in Mycobacterium tuberculosis. Mol Cell 81, 2887–2900 e2885

63. Mistry, J., Chuguransky, S., Williams, L., Qureshi, M., Salazar, G. A., Sonnhammer, E. L. L., Tosatto, S. C. E., Paladin, L., Raj, S., Richardson, L. J., Finn, R. D., and Bateman, A. (2021) Pfam: The protein families database in 2021. Nucleic Acids Res 49, D412–D419

64. Larkin, M. A., Blackshields, G., Brown, N. P., Chenna, R., McGettigan, P. A., McWilliam, H., Valentin, F., Wallace, I. M., Wilm, A., Lopez, R., Thompson, J. D., Gibson, T. J., and Higgins, D. G. (2007) Clustal W and Clustal X version 2.0. Bioinformatics 23, 2947–2948

65. Price, M. N., Dehal, P. S., and Arkin, A. P. (2010) FastTree 2 - approximately maximum-likelihood trees for large alignments. PLoS One 5, e9490

66. Balaban, M., Moshiri, N., Mai, U., Jia, X., and Mirarab, S. (2019) TreeCluster: Clustering biological sequences using phylogenetic trees. PLoS One 14, e0221068

67. Jumper, J., Evans, R., Pritzel, A., Green, T., Figurnov, M., Ronneberger, O., Tunyasuvunakool, K., Bates, R., Zidek, A., Potapenko, A., Bridgland, A., Meyer, C., Kohl, S. A. A., Ballard, A. J., Cowie, A., Romera-Paredes, B., Nikolov, S., Jain, R., Adler, J., Back, T., Petersen, S., Reiman, D., Clancy, E., Zielinski, M., Steinegger, M., Pacholska, M., Berghammer, T., Bodenstein, S., Silver, D., Vinyals, O., Senior, A. W., Kavukcuoglu, K., Kohli, P., and Hassabis, D. (2021) Highly accurate protein structure prediction with AlphaFold. Nature 596, 583–589

68. Waterhouse, A., Bertoni, M., Bienert, S., Studer, G., Tauriello, G., Gumienny, R., Heer, F. T., de Beer, T. A. P., Rempfer, C., Bordoli, L., Lepore, R., and Schwede, T. (2018) SWISS-MODEL: homology modelling of protein structures and complexes. Nucleic Acids Res 46, W296–W303

69. Abramson, J., Adler, J., Dunger, J., Evans, R., Green, T., Pritzel, A., Ronneberger, O., Willmore, L., Ballard, A. J., Bambrick, J., Bodenstein, S. W., Evans, D. A., Hung, C. C., O’Neill, M., Reiman, D., Tunyasuvunakool, K., Wu, Z., Zemgulyte, A., Arvaniti, E., Beattie, C., Bertolli, O., Bridgland, A., Cherepanov, A., Congreve, M., Cowen-Rivers, A. I., Cowie, A., Figurnov, M., Fuchs, F. B., Gladman, H., Jain, R., Khan, Y. A., Low, C. M. R., Perlin, K., Potapenko, A., Savy, P., Singh, S., Stecula, A., Thillaisundaram, A., Tong, C., Yakneen, S., Zhong, E. D., Zielinski, M., Zidek, A., Bapst, V., Kohli, P., Jaderberg, M., Hassabis, D., and Jumper, J. M. (2024) Accurate structure prediction of biomolecular interactions with AlphaFold 3. Nature 630, 493–500

70. Crooks, G. E., Hon, G., Chandonia, J. M., and Brenner, S. E. (2004) WebLogo: a sequence logo generator. Genome Res 14, 1188–1190

71. Robert, X., and Gouet, P. (2014) Deciphering key features in protein structures with the new ENDscript server. Nucleic Acids Res 42, W320–324

72. Sievers, F., and Higgins, D. G. (2018) Clustal Omega for making accurate alignments of many protein sequences. Protein Sci 27, 135–145

73. Letunic, I., and Bork, P. (2021) Interactive Tree Of Life (iTOL) v5: an online tool for phylogenetic tree display and annotation. Nucleic Acids Res 49, W293–W296

