## Supplementary Materials for "Aromatic Patch in WhiB-Like Transcription Factors Facilitates Primary Sigma Factor Interaction in *Mycobacterium tuberculosis*"

**This PDF file includes:**

SI Tables 1-2

SI Figures 1-12

SI References

### Supplementary Tables

### Table S1. Bacterial strains and plasmids used in this study.

| | Description | Reference |
|---|---|---|
| **Strains** | | |
| ***E. Coli*** | | |
| XL1-Blue | Host strain for routine cloning work | Stratagene |
| BL21-Gold (DE3) | Host strain for recombinant protein expression | Agilent Technology |
| ***M. Smegmatis (Msm)*** | | |
| Msm WT | *Mycobacterium smegmatis* $MC^2$ 155 (ATCC 700084), unmodified | ATCC |
| ***Plasmids for expression and purification of proteins from E. coli*** | | |
| pCDF-1b-6HisMtb$\sigma^A_{C170}$ | The DNA fragment encoding the C-terminal $\sigma^A$ containing the last 170 residues (Residues 359-528) was amplified from *Mycobacterium tuberculosis* (*Mtb*) H37Rv genomic DNA and inserted into the KpnI-XhoI site of pCDF-1b for expressing $His_6$-$\sigma^A_{C170}$ used in the *in vitro* pull-down assay with WhiB4 and WhiB5, respectively; $Spec^+$ | (1) |
| pCDF-1b-6HisMtb$\sigma^A_{C170}$-Strep | A modification of pCDF-1b- $His_6$Mtb$\sigma^A_{C170}$ by site directed mutagenesis for expressing $\sigma^A_{C170}$ with a 6His-tag in the N-terminus and a Strep-tag in the C-terminus ($His_6$- $\sigma^A_{C170}$-Strep) used in the *in vitro* pull-down assay with WhiB5; $Spec^+$ | This study |
| pCDF-1b- 6HisMtb$\sigma^A_{C170}$-H516A | A modification of pCDF-1b- $His_6$Mtb$\sigma^A_{C170}$ by site directed mutagenesis for expressing $His_6$- $\sigma^A_{C170}$ with the H516A mutation used in the *in vitro* pull-down assay with WhiB4 and WhiB5, respectively; $Spec^+$ | (1) |
| pET28-6His$\sigma^A_{C112}$-$\beta_{tip}$ | The gene encoding the C-terminal domain of $\sigma^A$ containing the last 112 residues (Residues 417-528, denoted $\sigma^A_{C112}$) was amplified from pCDF-1b-6HisMtb$\sigma^A_{C170}$ and fused to the β-flap-tip helix (Residues 815-829) with an artificial linker (GSSGSG). The resulting DNA fragment was inserted into the NcoI/XhoI site of pET28b for expressing $His_6$-$\sigma^A_{C112}$-$\beta_{tip}$ used in the ITC experiments; $Kan^R$ | (2) |
| pET28-Strep$\sigma^A_{C112}$-$\beta_{tip}$ | A modification of pET28-6His$\sigma^A_{C112}$-$\beta_{tip}$ by site-directed mutagenesis for expressing $\sigma^A_{C112}$-$\beta_{tip}$ with an N-terminal Strep-tag used for the pull-down assay for WhiB4 and WhiB6, respectively; $Kan^+$ | This study |
| pET28-Strep$\sigma^A_{C112}$-$\beta_{tip}$ -H516A | A modification of pET28-Strep$\sigma^A_{C112}$-$\beta_{tip}$ by site-directed mutagenesis for expressing Strep-tagged $\sigma^A_{C112}$-$\beta_{tip}$ with the H516A mutation used for the pull-down assay for WhiB4 and WhiB6; $Kan^+$ | This study |
| pET28-6His$\sigma^A_{C82}$-$\beta_{tip}$ | A modification of pET28-6His$\sigma^A_{C112}B_{tip}$ by site-directed mutagenesis for expressing the shorter chimera $His_6$-$\sigma^A_{C82}$-$\beta_{tip}$ This plasmid was used together with pET21-MtbWhiB6 for co-purification and crystallographic characterization of the WhiB6:$His_6$-$\sigma^A_{C82}$-$\beta_{tip}$ complex; $Kan^+$ | (2,3) |

| | | |
|---|---|---|
| pET21-MtbWhiB4 | The full-length *whiB4* gene was amplified from the *Mtb* genomic DNA and inserted into the NdeI/XhoI site of pET21 for expressing tagless MtbWhib4 used in the pull-down assay with $His_6$-Mtb$\sigma^{A}_{C170}$ (WT and mutant); $Amp^{+}$ | This study |
| pET21-MtbWhiB4-His6 | A modification of pET21-WhiB4 by site-directed mutagenesis for expressing WhiB4-$His_6$ used in the pull-down assay with Strep-$\sigma^{A}_{C112}B_{tip}$; $Amp^{+}$ | This study |
| pET21-MtbWhiB4-His6 mutant (L45A, F46A) | A modification of pET21-MtbWhiB4-His6 by site-directed mutagenesis for expressing WhiB4-$His_6$ with a single mutation of either L45A or F46A used in the pull-down assay with Strep-$\sigma^{A}_{C112}B_{tip}$; $Amp^{+}$ | This study |
| pET21-MtbWhiB5 | The full-length *whiB5* gene (Rv0022c) was amplified from the *Mtb* genomic DNA and inserted into the NdeI/XhoI site of pET21 for expressing tagless *Mtb* WhiB5 used in the pull-down assay with $His_6$-Mtb$\sigma^{A}_{C170}$ (WT and mutant); $Amp^{+}$ | This study |
| pET21-MtbWhiB5 mutant (W12A, F13A) | A modification of pET21-WhiB5 by site-directed mutagenesis for expressing tagless WhiB5 with a single mutation of either W12A or F13A used in the pull-down assay with $His_6$-Mtb$\sigma^{A}_{C170}$; $Amp^{+}$ | This study |
| pET21-MtbWhiB6 | The DNA fragment encoding 116-aa *Mtb* WhiB6 reported in the MycoBrowser portal (Rv3862c) was amplified from the *Mtb* genomic DNA and inserted into the NdeI/XhoI site of pET21 for expressing the tagless 116-aa *Mtb* WhiB6 for co-purification and crystallographic characterization of the WhiB6:$His_6$-$\sigma^{A}_{C82}$-$\beta_{tip}$ complex; $Amp^{+}$ | This study |
| pET21-MtbWhiB6Short-6His | A modification of pET21-MtbWhiB6 by site-directed mutation to remove the 60 bp of the 5'-*whiB6* gene in pET21-MtbWhiB6 based on the refined the redefined open reading frame of WhiB6 (96 aa) in the study by Shell, *et al.* (3). The resulting plasmid was used for expressing a C-terminal $His_6$ tagged WhiB6 used in the pull-down assay with Strep-$\sigma^{A}_{C112}$-$\beta_{tip}$ (WT and mutant); $Amp^{+}$ | This study |
| pET21-MtbWhiB6Short-6His (W21A, T22A) | A modification of pET21-MtbWhiB6Short-6His by site-directed mutagenesis for expressing the $His_6$-WhiB6 mutant carrying a single mutation of either W21A or T22A used in the pull-down assay with Strep-$\sigma^{A}_{C112}$-$\beta_{tip}$; $Amp^{+}$ | This study |
| pET21-A0A0K2FNL9-His6 | The DNA fragment encoding a Wbl homolog (Uniprot ID: A0A0K2FNL9) from *Mycobacterium phage Lolly9* was synthesized by GenScript and inserted into the NdeI/XhoI site of pET21 for expressing A0A0K2FNL9-$His_6$ and used in the pull-down assay with Strep-$\sigma^{A}_{C112}$-$\beta_{tip}$; $Amp^{+}$ | This study |
| pET21-A0A0K2FNL9-His6 (F17A, F18A, W4A-F58A, W4A-F17A-F58A) | A modification of pET21-A0A0K2FNL9-$His_6$ by site-directed mutagenesis for expressing the A0A0K2FNL9-$His_6$ mutant carrying either a single mutation (F17A or F18A), double mutation (W4A-F58A), or triple mutation (W4A-F17A-F58A), and used in the pull-down assay with Strep-$\sigma^{A}_{C112}$-$\beta_{tip}$; $Amp^{+}$ | This study |
| pET21-Q857R7-His6 | The DNA fragment encoding a Wbl homolog (Uniprot ID: Q857R7) from the *Mycobacterium phage CJW1* was synthesized by GenScript and inserted into the NdeI/XhoI site of pET21 for expressing Q857R7-$His_6$ and used in the pull-down assay with Strep-$\sigma^{A}_{C112}$-$\beta_{tip}$; $Amp^{+}$ | This study |

| | | |
|---|---|---|
| pET21-Q857R7-His6 (F20A, F21A) | A modification of pET21-Q857R7-$His_6$ by site-directed mutagenesis for expressing the Q857R7-$His_6$ mutant carrying a single mutation of either F20A or F21A and used in the pull-down assay with Strep-$\sigma^A_{C112}$-$\beta_{tip}$; $Amp^+$ | This study |
| pET21- A0A3G3M9S7-His6 | The DNA fragment encoding a Wbl homolog (Uniprot ID: A0A3G3M9S7) from *Gordonia phage Octobien14* was synthesized by GenScript and inserted into the NdeI/XhoI site of pET21 for expressing A0A3G3M9S7-$His_6$ used in the pull-down assay with Strep-$\sigma^A_{C112}$-$\beta_{tip}$; $Amp^+$ | This study |
| pET21- A0A3G3M9S7-His6 (W12A, Y26A) | A modification of pET21-A0A3G3M9S7-$His_6$ by site-directed mutagenesis for expressing the A0A3G3M9S7-$His_6$ mutant carrying a single mutation of either W12A or Y26A and used in the pull-down assay with Strep-$\sigma^A_{C112}$-$\beta_{tip}$; $Amp^+$ | This study |
| pETDuet-6His-SUMO-MtbWhiB1 | The full length *whiB1* gene (Rv3219) was amplified from *M. tuberculosis* H37Rv genomic DNA and inserted into the NdeI/XhoI site of a modified pETDuet-SUMO for expressing $His_6$-SUMO-WhiB1 that was used in the ITC assay; $Amp^R$ | (1) |
| pETDuet-6His-SUMO-MtbWhiB2 | The full length *whiB2* gene (Rv3260c) was amplified from *M. tuberculosis* H37Rv genomic DNA and inserted into the NdeI/XhoI site of a modified pETDuet-SUMO for expressing $His_6$-SUMO-WhiB2 that was used in the ITC assay; $Amp^R$ | This study |
| pETDuet-6His-SUMO-MtbWhiB5 | The full length *whiB5* gene (Rv0022c) was amplified from pET21-MtbWhiB5 and inserted into the NdeI/XhoI site of a modified pETDuet-SUMO for expressing $His_6$-SUMO-WhiB5 that was used in the ITC assay; $Amp^R$ | This study |
| ***Plasmids for Mycobacterium Protein fragmentation (M-PFC) assay in Msm*** | | |
| pUAB100 | The plasmid encodes the *Saccharomyces cerevisiae* leucine-zipper sequence homodimerization domain (GCN4) that is fused to the N terminus of dihydrofolate reductase fragment ($mDHFR_{F[1,2]}$) with a $Gly_{10}$ linker GCN4-$Gly_{10}$- $mDHFR_{F[1,2]}$ under the control of the *hsp60* promoter. This plasmid was co-transformed into *Msm* with pUAB200 and used as a positive control in the M-PFC assay; $Hyg^+$ | (4) |
| pUAB200 | The plasmid encodes the *Saccharomyces cerevisiae* leucine-zipper sequence homodimerization domain (GCN4) that is fused to the N terminus of dihydrofolate reductase fragment ($mDHFR_{F[3]}$) with a $Gly_{10}$ linker GCN4-$Gly_{10}$- $mDHFR_{F[3]}$ under the control of the *hsp60* promoter. This plasmid was co-transformed into *Msm* with pUAB100 and used as a positive control in the M-PFC assay; $Kan^+$ | (4) |

| | | |
|---|---|---|
| pUAB300 | The plasmid encodes the dihydrofolate reductase fragment ($mDHFR_{F[1,2]}$) with a $Gly_{10}$ linker under the control of the *hsp60* promoter. This plasmid was co-transformed into *Msm* with pUAB400 and used as a negative control in the M-PFC assay; $Hyg^+$ | (4) |
| pUAB400 | The plasmid encodes the dihydrofolate reductase fragment ($mDHFR_{F[3]}$) with a $Gly_{10}$ linker under the control of the *hsp60* promoter. This plasmid was co-transformed into *Msm* with pUAB300 and used as a negative control in the M-PFC assay; $Kan^+$ | (4) |
| pUAB100-WhiB5 | The DNA fragment encoding GCN4 in pUAB100 was removed and replaced by the gene encoding the full-length *Mtb* WhiB5 amplified from pET21-MtbWhiB5. The resulting plasmid was used to express WhiB5-$Gly_{10}$-$mDHFR_{F[1,2]}$ under the control of the *hsp60* promoter. This plasmid was co-transformed into *Msm* with pUAB200-SigAc82btip to test the interaction between WhiB5 and $\sigma^A{}_4$ in the M-PFC assay; $Hyg^+$ | This study |
| pUAB100-WhiB5 mutants (W12A, F13A) | Modified from the pUAB100-WhiB5 plasmid by site-directed mutagenesis to express WhiB5-$Gly_{10}$-$mDHFR_{F[1,2]}$ with a single mutation of either W12A or F13A under the control of the *hsp60* promoter. This plasmid was co-transformed into *Msm* with pUAB200-SigAc82btip to test the interaction between the WhiB5 mutants and $\sigma^A{}_4$ in the M-PFC assay; $Hyg^+$ | This study |
| pUAB200-$\sigma^A{}_{C82}$-$\beta_{tip}$ | The DNA fragment encoding GCN4 in pUAB200 was removed and replaced by the DNA fragment encoding the $\sigma^A{}_{C82}$-$\beta_{tip}$ chimera amplified from pET28-$\sigma^A{}_{C82}$-$\beta_{tip}$. The resulting plasmid was used to express $\sigma^A{}_{C82}B_{tip}$-$Gly_{10}$- $mDHFR_{F[3]}$ under the control of the *hsp60* promoter. This plasmid was co-transformed into *Msm* with pUAB100-WhiB5 (either wildtype or mutants) to test the interaction between WhiB5 and $\sigma^A{}_4$ in the M-PFC assay; $Kan^+$ | This study |

**Supplementary Table 2. Data collection and refinement statistics.**

| Proteins | WhiB6:$\sigma^{A}{}_{c82}$-$\beta_{tip}$ (phasing) | WhiB6:$\sigma^{A}{}_{c82}$-$\beta_{tip}$ |
|---|---|---|
| **Data collection**[a] | | |
| Space group | $C222_1$ | $C222_1$ |
| Cell dimensions | | |
| *a*, *b*, *c* (Å) | 63.5, 67.0, 159.2 | 63.0, 66.5, 158.8 |
| α, β, γ (°) | 90.0, 90.0, 90.0 | 90.0, 90.0, 90.0 |
| Wavelength (Å) | 1.7365 | 0.9795 |
| Resolution (Å) | 50-2.30 (2.34-2.30) | 50-1.8 (1.864-1.8) |
| $R_{merge}$ | 0.129 (1.610) | 0.067 (1.59) |
| $I / \sigma I$ | 23.6 (1.08) | 18.0 (1.5) |
| Completeness (%) | 96.4 (91.4) | 97.4 (97.5) |
| Multiplicity | 21.3 (11.6) | 12.7 (13.6) |
| No. unique reflections | | 30736 (3011) |
| $CC_{1/2}$(%) | 100.0 (63.5) | 99.9 (80.9) |
| **Refinement** | | |
| Resolution (Å) | | 50-1.8 (1.864-1.8) |
| No. molecules per asymmetric unit | | 2 |
| $R_{work}/R_{free}$ | | 0.213/0.232 |
| Included residue No. | | |
| WhiB6 | | 8–91; 9-90[b] |
| $\sigma^{A}{}_{4}$ | | 453–528; 454–528 |
| $\beta_{tip}$ | | 815–826; 815–824 |
| No. atoms | | 2951 |
| Macromolecules | | 2798 |
| Ligand | | 16 |
| Water | | 137 |
| Avg. *B*-factors (Å$^2$) | | 54.3 |
| Macromolecules | | 54.6 |
| Ligand | | 42.7 |

| | | |
|---|---|---|
| Water | | 48.5 |
| r.m.s deviations | | |
| Bond lengths (Å) | | 0.01 |
| Bond angles (°) | | 1.21 |
| Ramachandran statistics | | |
| Favored regions (%) | | 97.42 |
| Allowed regions (%) | | 2.58 |
| Outliers (%) | | 0 |
| PDB code | | 8D5V |

[a]The highest resolution shell statistics are shown in parentheses.

[b]The two sets of the residues are for each of the two complexes in the asymmetric unit.

**Supplemental Figures**

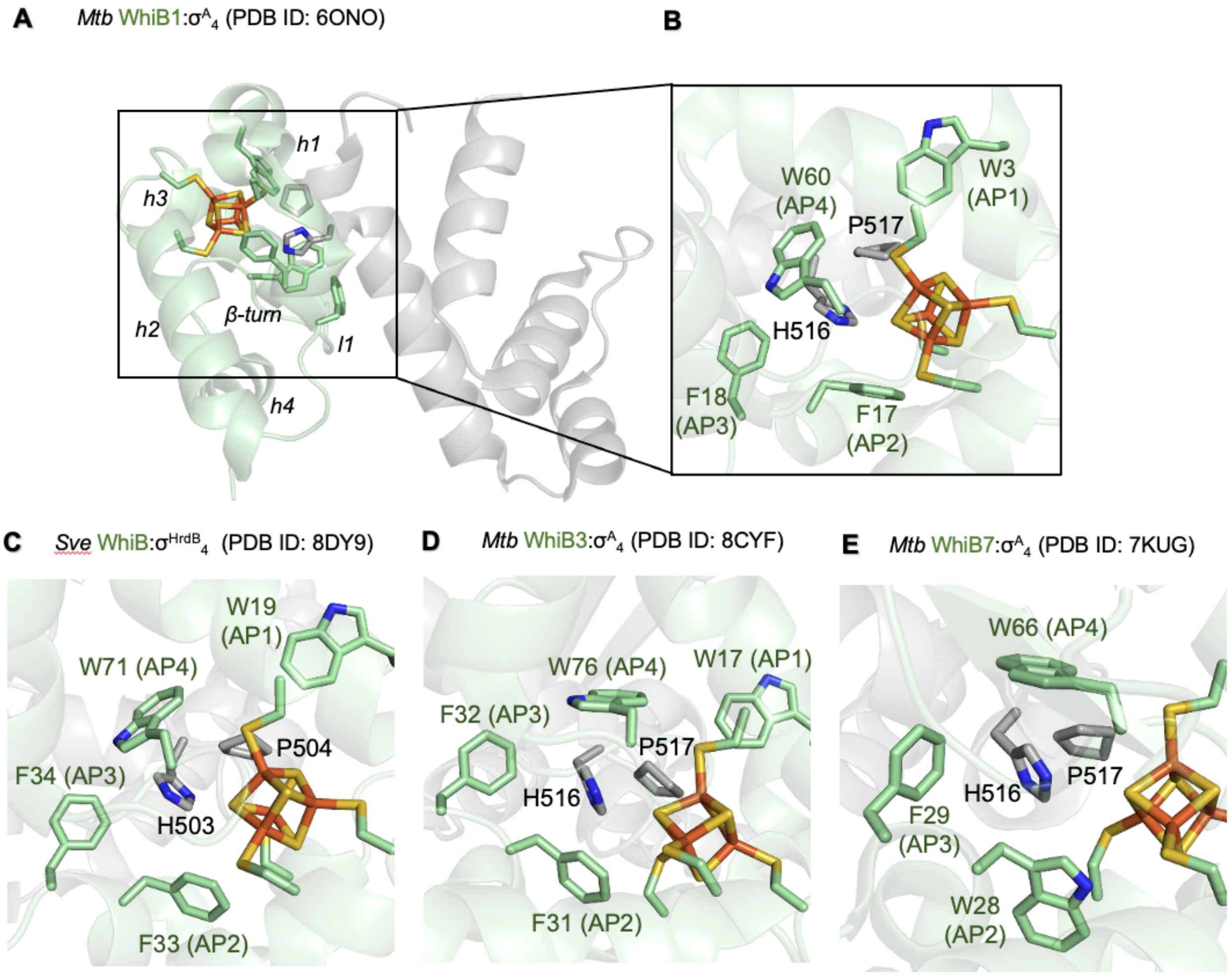


**Supplementary Figure 1. Highlights of the aromatic patch in the previously characterized $\sigma^A{}_4$-bound Wbls from *Mtb* and from *Streptomyces venezuelae* (*Sve*).** (A) Overall structure of the $\sigma^A{}_4$-bound *Mtb* WhiB1 and (B) Zoom-In view of the aromatic patch at WhiB1:$\sigma^A{}_4$ interface within the Fe-S cluster binding pocket. (C-E) the Zoom-In view of the aromatic patch at the Wbl:$\sigma^A{}_4$ interface within the Fe-S cluster binding pocket of *Sve* WhiB:$\sigma^A{}_4$, *Mtb* WhiB3:$\sigma^A{}_4$ and *Mtb* WhiB7:$\sigma^A{}_4$, respectively. *Sve* WhiB is a homolog of *Mtb* WhiB2. In all structures, Wbl proteins are colored pale green and $\sigma^A{}_4$ in grey. The [4Fe-4S] cluster, the two $\sigma^A$ residues (H516-P517 in *Mtb* $\sigma^A$; H503-P504 in *Sve* $\sigma^A$ [also known as $\sigma^{HrdB}$]) and the aromatic patch of Wbls at the molecular interface between $\sigma^A{}_4$ and Wbls are shown in sticks. Fe atoms are colored orange, S in yellow, O in red, and N in blue. AP1-4: the residues in the aromatic patch residues 1-4 (AP1-4) corresponding to the W3, F17, F18 and W60, respectively, in WhiB1.

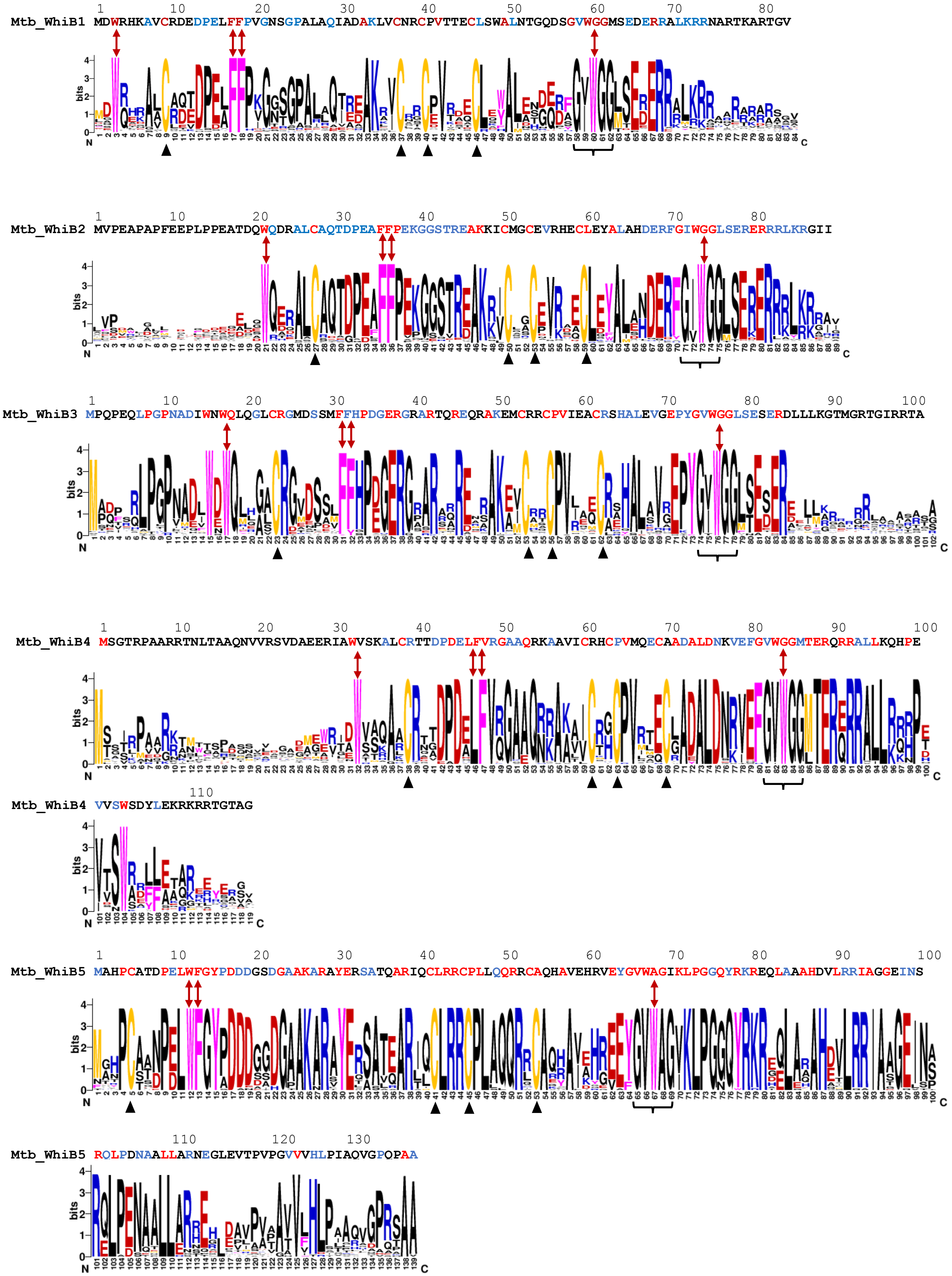
Mtb_WhiB1 MDWRHKAVCRDEDPELFFPVGNSGPALAQIADAKLVCNRCPVTTECLSWALNTGQDSGVWGGMSEDERRALKRRNARTKARTGV
Mtb_WhiB2 MVPEAPAPFEEPLPPEATDQWQDRALCAQTDPEAFFPEKGGSTREAKKICMGCEVRHECLEYALAHDERFGIWGGLSERERRRLKRGII
Mtb_WhiB3 MPQPEQLPGPNADIWNWQLQGLCRGMDSSMFFHPDGERGRARTQREQRAKEMCRRCPVIEACRSHALEVGEPYGVWGGLSESERDLLLKGTMGRTGIRRTA
Mtb_WhiB4 MSGTRPAARRTNLTAAQNVVRSVDAEERIAWVSKALCRTTDPDELFVRGAAQRKAAVICRHCPVMQECAADALDNKVEFGVWGGMTERQRRALLKQHPE
Mtb_WhiB4 VVSWSDYLEKRKRRTGTAG
Mtb_WhiB5 MAHPCATDPELWFGYPDDDGSDGAAKARAYERSATQARIQCLRRCPLLQQRRCAQHAVEHRVEYGVWAGIKLPGGQYRKREQLAAAHDVLRRIAGGEINS
Mtb_WhiB5 RQLPDNAALLARNEGLEVTPVPGVVVHLPIAQVGPQPAA

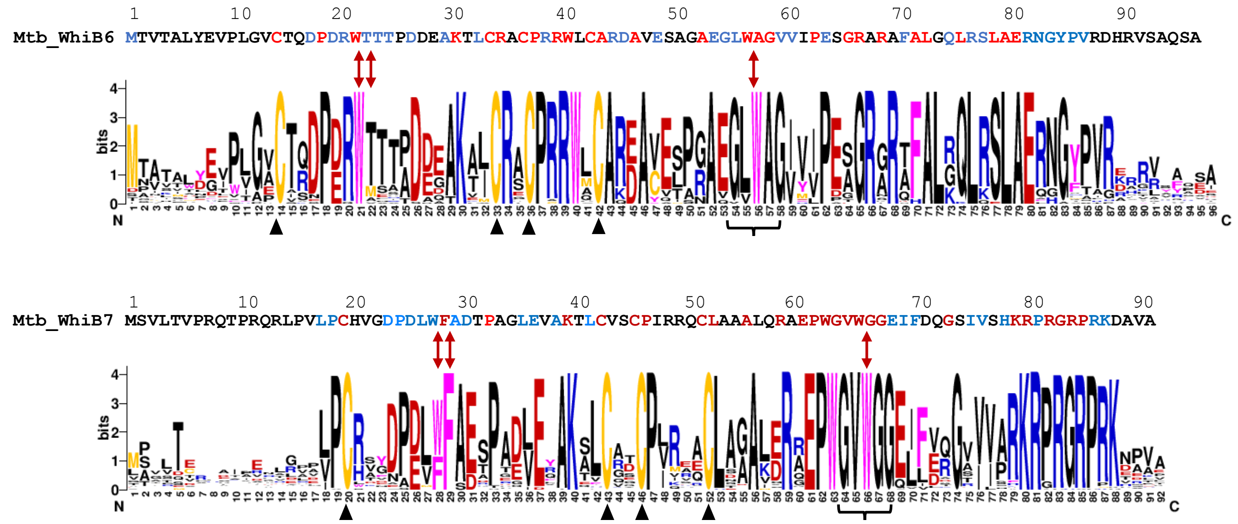


**Supplementary Figure 2. Sequence logos of the selected Wbl subfamilies proteins.** The protein sequences of *Mtb* Wbls are followed by the sequence logos of the members in the corresponding subfamilies. The invariant residues and the highly conserved residues (varied residues with a similar polarity) are highlighted in red and blue, respectively, in the *Mtb* Wbl protein sequences. The numbering of the residues in the sequence logos follows the *Mtb* Wbl proteins, with the alignment gaps in the sequence logos removed for clarity. The corresponding residues of the aromatic patch are highlighted by the red arrows, and the four conserved Cys are indicated by the black arrows. The sequence logos for the subfamilies of WhiB1, WhiB3, and WhiB7 are modified from the previous reports (2,5). For the WhiB2 subfamily, *Mtb* WhiB2 was aligned against WhiB2 was aligned against 99 orthologs in Actinobacteria, including *Streptomycineae, S. albus, S. avermitilis, S. clavuligerus, S. griseoflavus Tu4000, S. griseus subsp. griseus, S. hygroscopicus, S. lividans TK24, S. pristinaespiralis, S. scabiei, S. venezuelae, S. viridochromogenes DSM 40736, S. sviceus ATCC 29083, S. filamentosus NRRL 11379; Kitasatospora setae KM-60(2,5)4; Catenulispora acidiphila DSM 44928; Salinispora tropica CNB-440; Micromonospora, M. sp. L5, M. aurantiaca, M. Llam0; Saccharomonospora viridis DSM 43017; Saccharopolyspora erythraea; Actinosynnema mirum DSM 43827; Amycolatopsis mediterranei U32; Thermobispora bispora; Streptosporangium roseum DSM 43021; Thermomonospora curvata; Thermobifida fusca; Nocardiopsis dassonvillei subsp. dassonvillei DSM 43111; Acidothermus cellulolyticus 11B; Frankia, F. alni ACN14a, F. sp., F. sp. CcI6; Geodermatophilus obscurus DSM 43160; Nakamurella multipartita DSM 44233; Gordonia bronchialis DSM 43247; Nocardia farcinica IFM 10152; Segniliparus rotundus DSM 44985; Tsukamurella paurometabola DSM 20162; Rhodococcus, R. erythropolis PR4, R. jostii RHA1, R. opacus B4, R. hoagii 103S; Mycobacteria, M. vanbaalenii PYR-1, M. ulcerans Agy99, M. smegmatis str. MC2 155, M. marinum M, M. leprae, M. gilvum PYR-GCK, M. abscessus, M. avium subsp. paratuberculosis K-10, M. avium, M. tuberculosis variant bovis, M. sp. PAM1, M. gilvum; Corynebacterium, C. urealyticum, C. pseudotuberculosis, C. kroppenstedtii DSM 44385, C. jeikeium, C. glutamicum ATCC 13032 2, C. efficiens, C. diphtheriae, C. aurimucosum ATCC 700975; Nocardioides sp.; Kribbella flavida DSM 17836; Propionibacterium, P. freudenreichii, P. acnes; Kineococcus radiotolerans; Beutenbergia cavernae; Cellulomonas flavigena DSM 20109; Brachybacterium faecium DSM*

*4810; Kytococcus sedentarius; Intrasporangium calvum; Jonesia denitrificans; Clavibacter michiganensis subsp. nebraskensis; Leifsonia xyli subsp. xyli str. CTCB07; Microbacterium thalassium; Arthrobacter, A. sp. FB24, A. arilaitensis, A. arilaitensis, A. aurescens TC1, A. chlorophenolicus, A. phenanthrenivorans; Kocuria rhizophila DC2201; Micrococcus luteus NCTC 2665; Renibacterium salmoninarum ATCC 33209; Rothia, R. dentocariosa, R. mucilaginosa; Xylanimonas allomyrinae; Sanguibacter keddieii DSM 10542; Tropheryma whipplei str. Twist; Mobiluncus curtisii ATCC 43063; Arcanobacterium haemolyticum; Bifidobacterium, B. adolescentis, B. bifidum, B. dentium, B. longum, B. longum NCC2705, B. crudilactis; Acidimicrobium ferrooxidans DSM10331.* For the WhiB4 subfamily, *Mtb* WhiB4 was aligned against 45 orthologs in Actinobacteria, including *Streptomycineae, S. lividans, S. viridochromogenes, S. scabiei, , S. hygroscopicus, S. sviceus, S. avermitilis, S. griseoflavus, S. venezuelae, S. griseus, S. pristinaespiralis, S. albus, S. clavuligerus; Kitasatospora setae; Stackebrandtia nassauensis; Micromonospora aurantiaca; Saccharomonospora viridis; Amycolatopsis mediterranei; Thermobispora bispora; Thermomonospora curvata; Thermobifida fusca; Acidothermus cellulolyticus; Gordonia bronchialis; Nocardia farcinica; Tsukamurella paurometabola; Rhodococcus, R. jostii, R. erythropolis; Mycobacterium, M. vanbaalenii, M. ulcerans, M. gilvum, M. sp. MCS, M. sp. KMS, M. sp. JLS, M. smegmatis, M. marinum, M. leprae, M. bovis; Corynebacterium, C. jeikeium, C. urealyticum, C. pseudotuberculosis, C. glutamicum, C. efficiens, C. diphtheriae, C. aurimucosum; Kineococcus radiotolerans; Intrasporangium calvum.* For the WhiB5 subfamily, *Mtb* WhiB5 was aligned against 28 orthologs in Actinobacteria, including *Mycobacterium*, *M. sp. ACS4331, M. africanum, M. angelicum, M. bovis, M. decipiens, M. helveticum, M. kansasii, M. kubicae, M. lacus, M. marinum, M. pseudokansasii, M. simiae, M. confluentis, M. koreensis, M. intracellulare, M. nebraskense, M. parmense, M. sp. Marseille-P9652, M. sp. 852002-50816 SCH5313054-b, M. sp. 1100029.7, M. shinjukuense, M. riyadhense, M. botniense, M. palauense, M. icosiumassiliensis, M. sp. MS1601, M. sp. 1274756.6, M. sarraceniae, M. chitae, M. fallax.* For the WhiB6 subfamily, *Mtb* WhiB6 was aligned against 35 orthologs in mycobacteria, including *Mycobacterium, M. sp. ACS4331, M. sp. MCS, M. JLS, M. abscessus, M. africanum, M. angelicum, M. bovis, M. decipiens, M. helveticum, M. kansasii, M. kubicae, M. lacus, M. marinum, M. mungi, M. pseudokansasii, M. simiae, M. acapulense, M. agri, M. vanbaalenii, M. aurum, M. arabiense, M. celeriflavum, M. confluentis, M. diernhoferi, M. flavescens, M. frederiksbergense, M. gilvum, M. iranicum, M. koreensis, M. madagascariense, M. obuense, M. parafortuitum, M. smegmatis, M. septicum, M. setense.*

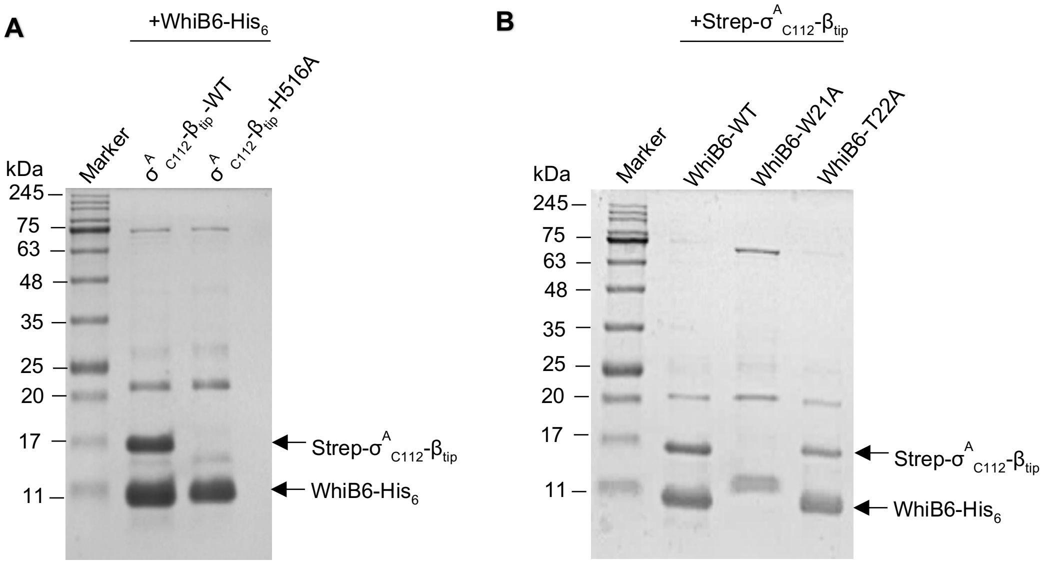


**Supplementary Figure 3. The uncropped SDS PAGE images of the pull-down samples shown in Fig. 1E and F, respectively.**

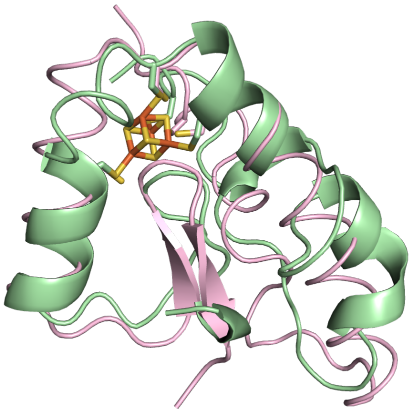


**Supplementary Figure 4. Comparison of the homologous model of WhiB5:σ$^A_4$ by SWISS-Model (pale green) with that of the *ab initio* model predicted AlphaFold3 (pink).** In both structures, σ$^A_4$ in the complexes is not shown for clarity. The TM score for the AlphaFold model is 0.67 for WhiB5.

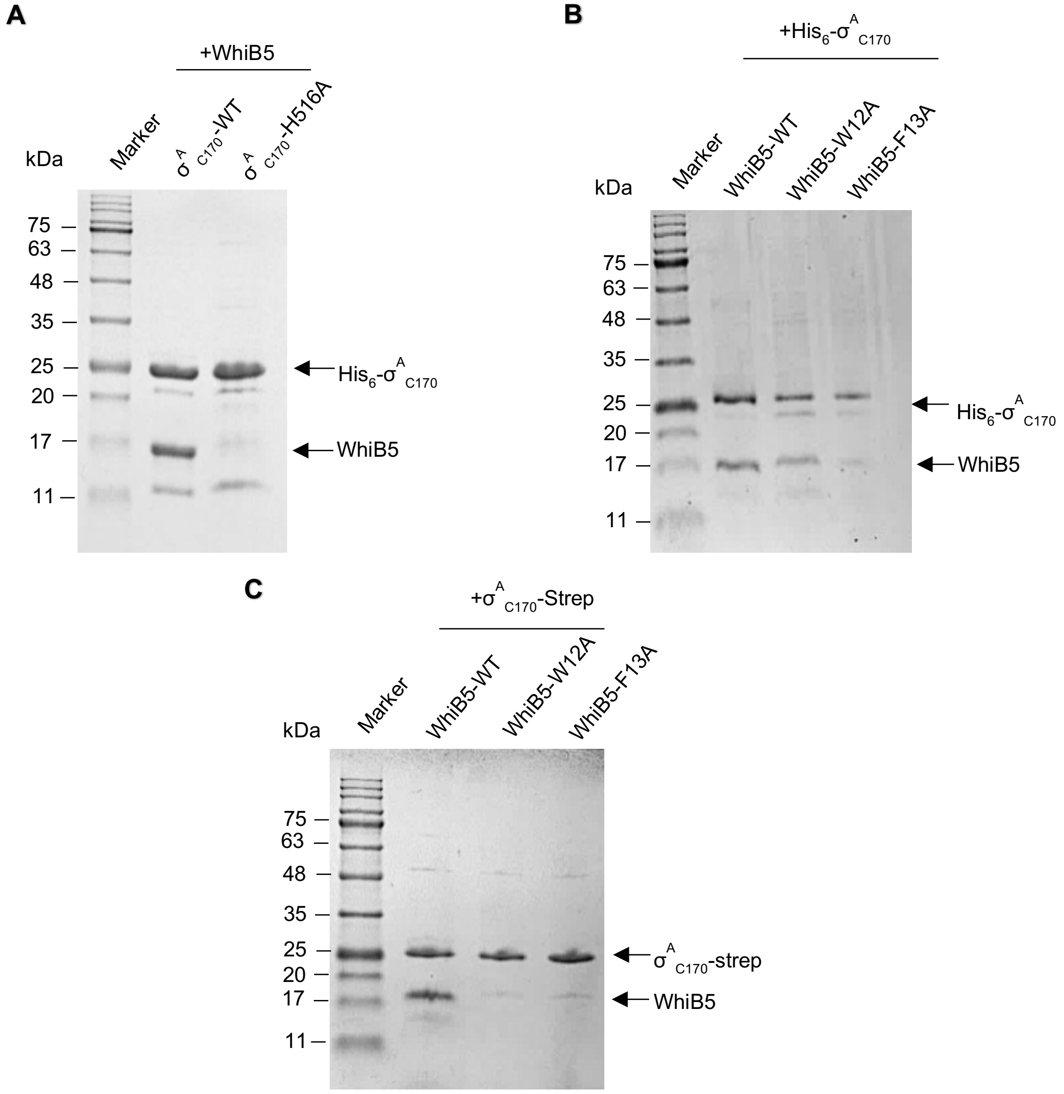


**Supplementary Figure 5. The uncropped SDS PAGE images of the pull-down samples shown in Fig. 2A, C and D, respectively.**

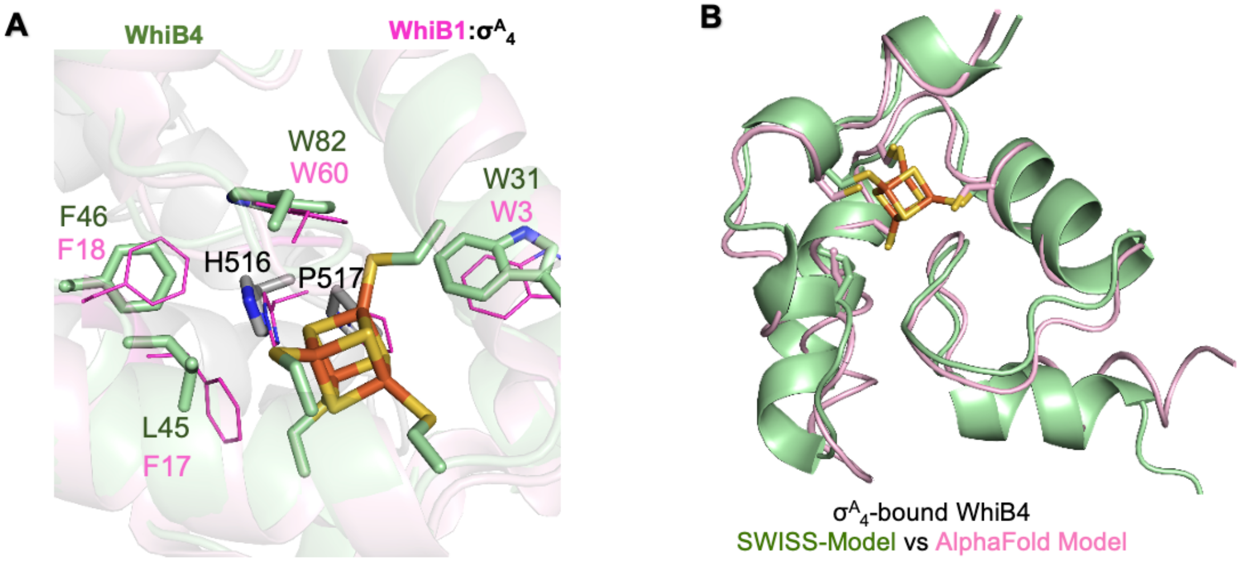


**Supplementary Figure 6. Structural models of the WhiB4:σ$^{A}_{4}$ complex.** (A) Zoom-in view of the homologous model of the σ$^{A}_{4}$-bound WhiB4 by SWISS-Model (pale green) at the WhiB4:σ$^{A}_{4}$ interaction interface around the [4Fe-4S] cluster binding pocket, overlaid with WhiB1:σ$^{A}_{4}$ (PDB ID:6ONO, pink). The [4Fe-4S] clusters and the H516-P517 pair in WhiB1:σ$^{A}_{4}$, and the residues corresponding to the aromatic patch residues AP1-4 in the Wbls are shown in sticks. Fe atoms are colored orange, S in yellow, O in red and N in blue. (B) Comparison of the homologous model of the σ$^{A}_{4}$-bound WhiB4 (pale green) with that of the *ab initio* model predicted by AlphaFold3 (pink). The TM score for the AlphaFold model is 0.76 for WhiB4. σ$^{A}_{4}$ in the complex model is not shown for clarity.

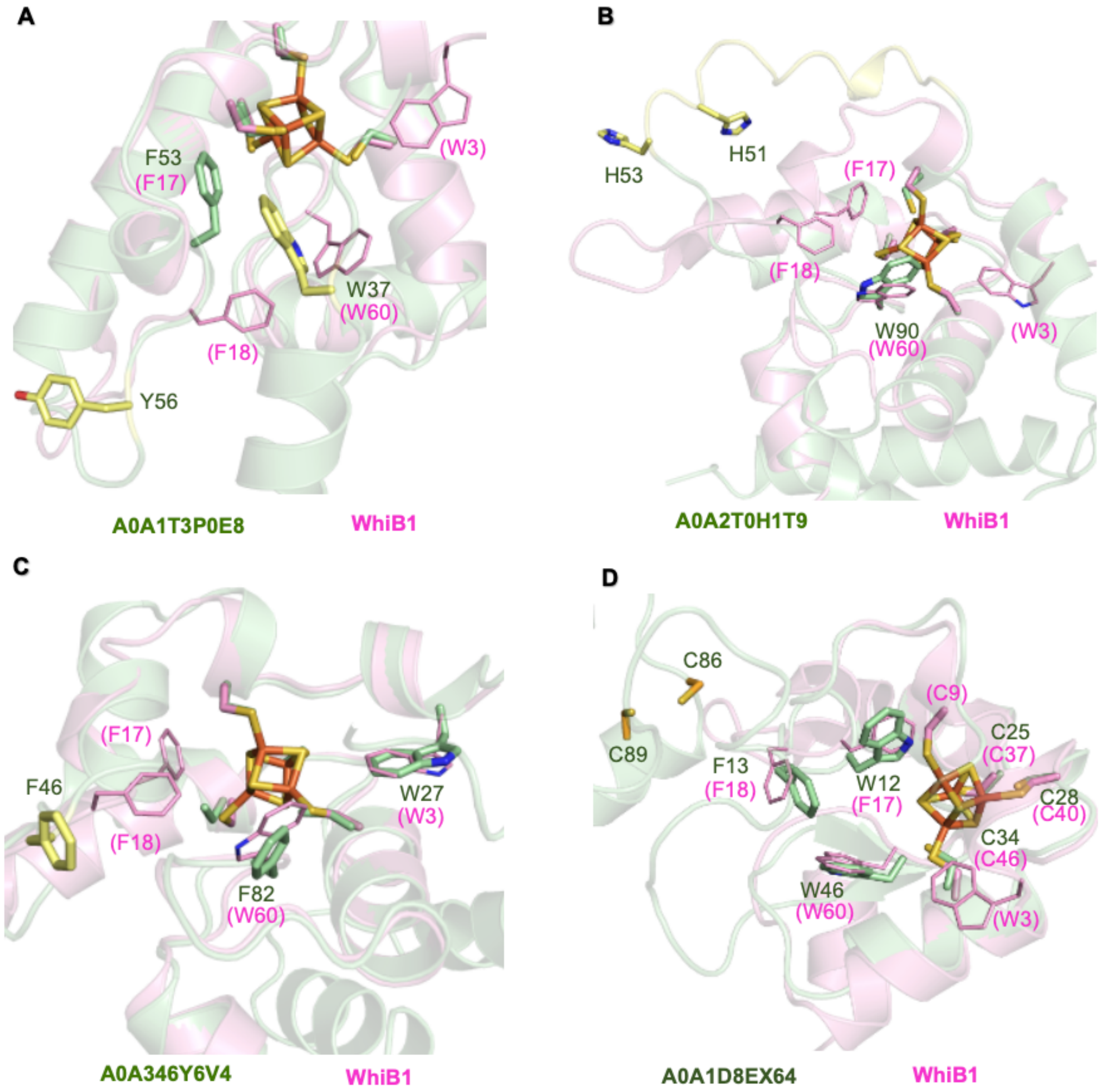


**Supplementary Figure 8. Comparison of $\sigma^{A}_{4}$-bound *Mtb* WhiB1 with the AlphaFold models of the Wbl outliers.** (A-C) The Wbls outliers lacking the aromatic patch, defined by at least two aromatic residues (either Trp, Phe, Tyr or His) corresponding to AP1-4 (W3, F17, F18 and W60 in WhiB1) and at least one corresponding to AP2-3 (F17 and F18 in WhiB1), including (A) A0A1T3P0E8 from *Embleya scabrispora* in the No. 7 Wbl subfamily, (B) A0A2T0H1T9 from *Actinopolyspora mortivallis* in the No. 21 Wbl subfamily, (C) A0A346Y6V4 from *Euzebya pacifica* in the No. 21 Wbl subfamily. (D) The Wbl outlier, A0A1D8EX64 from *Mycobacterium phage Tortellini49* in the No.17 Wbl subfamily, with only three Cys found at the expected [4Fe-4S] cluster binding pocket and missing a Cys residue corresponding to C9 in WhiB1. In all the structures, WhiB1 is colored in pink and the Wbl outliers are in pale green. The [4Fe-4S] cluster, the Cys ligands and the residues in the aromatic patch of WhiB1 are highlighted as references. The Wbl residues corresponding to the aromatic patch in WhiB1 are highlighted in sticks with the C atoms colored pale green. The other cysteines and aromatic residues outside the [4Fe-4S] cluster binding site are highlighted in sticks with the C atoms in yellow. Fe atoms are colored orange, S in yellow, O in red, and N in blue. $\sigma^{A}{}_{4}$ is not shown for clarity. The TM scores of the AlphaFold3 models of these Wbls:$\sigma^{A}{}_{4}$-$\beta_{tip}$ complexes range from high to low confidence using a TM score of 0.5 as the threshold (6): 0.83 for A0A1D8EX64 (D), 0.59 for A0A346Y6V4 (B), 0.51 for A0A2T0H1T9 (A), and 0.42 for A0A1T3P0E8 (C).

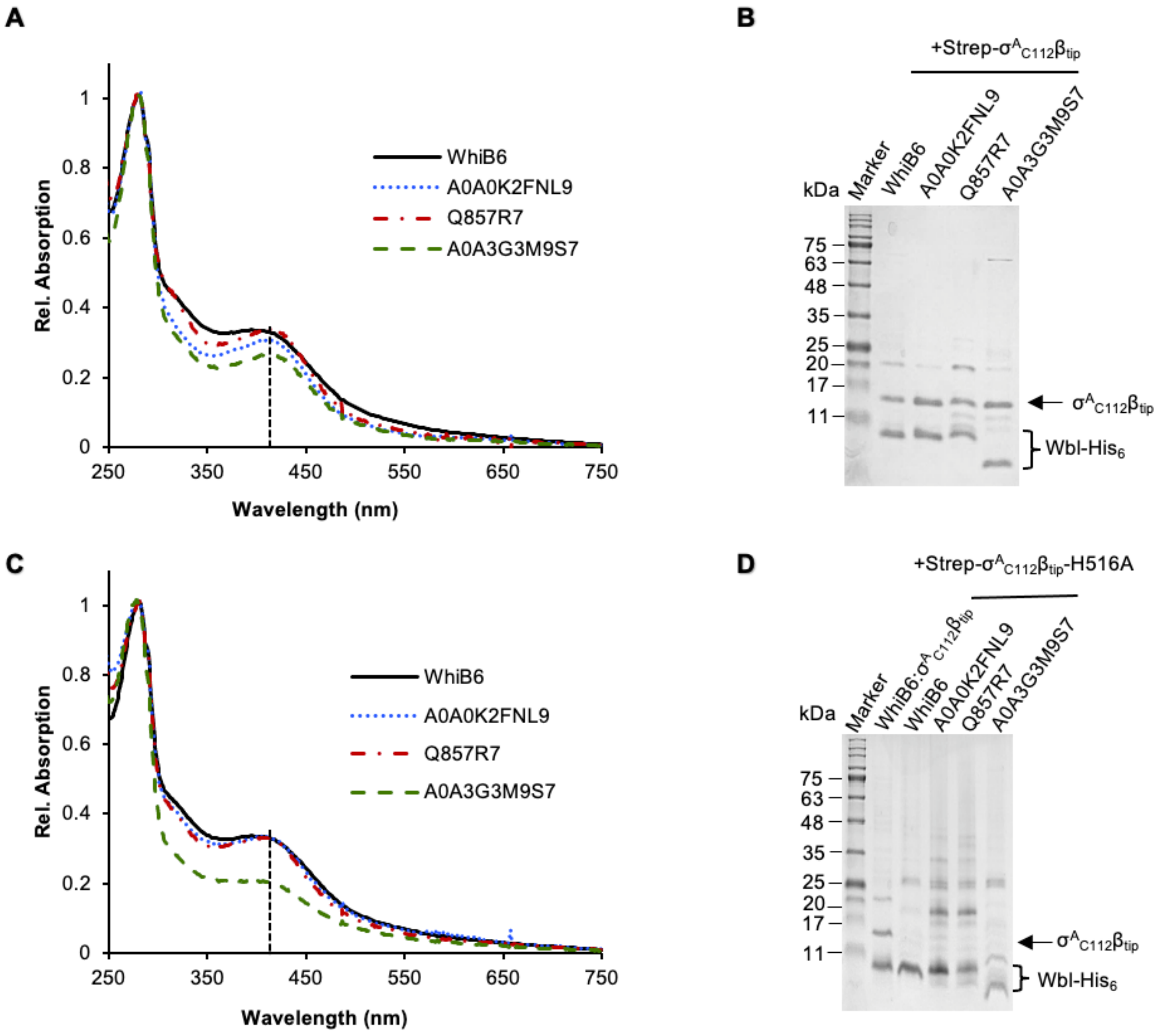


**Supplementary Figure 9. Interactions between the Wbls from actinobacteriophages with *Mtb* $\sigma^A_4$ in a $\sigma^A_4$-H516 dependent manner by pull-down assays.** The three phage Wbls are referred to by their UniProt IDs: A0A0K2FNL9 from *Mycobacterium phage Lolly9* belonging to the No. 19 Wbl subfamily; Q857R7 from *Mycobacterium virus CJW1* in the WhiB1 subfamily; and A0A3G3M9S7 from *Gordonia phage Octobien14* in the No. 20 Wbl subfamily (Fig. 5, Supplementary DataSet 1). (A) and (B), UV-Visible spectra and SDS-PAGE analyses of the samples from co-expression and affinity purification of $\sigma^A_4$ and Wbls. (C) and (D), UV-Visible spectra and SDS-PAGE analyses of the samples from co-expression and purification of $\sigma^A_4$-H516A mutant and Wbls-$His_6$ as indicated. In both assays, Wbl-$His_6$ was used as the bait. WhiB6:$\sigma^A_4$ was used as the reference. The absorption spectra were normalized to the absorbance at 280 nm. The intensity of the absorption peak around 410 nm, marked with a dashed line, is indicative of the occupancy of the [4Fe-4S] cluster in the samples. The reduced intensity of the 410-nm peak for the A0A3G3M9S7 sample in Panel (C) compared to that in Panel (A) indicates a cluster loss without the protection of $\sigma^A_4$, while the other two phage Wbls show no significant change.

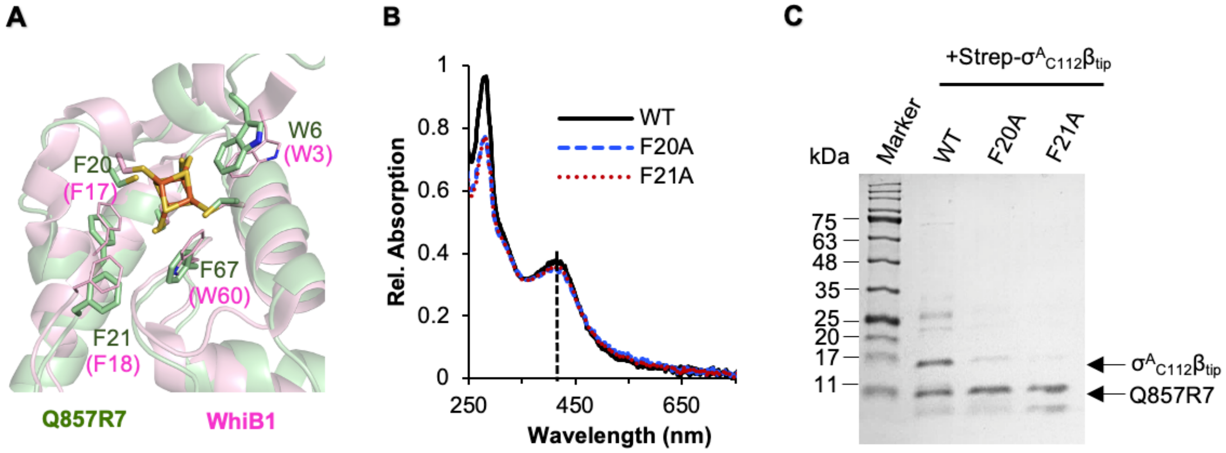


**Supplementary Figure 10. Effect of aromatic patch residues in the Wbl (UniProt ID: Q857R7) from *Mycobacterium virus CJW1* on $\sigma^A{}_4$ binding.** (A) Comparison of the AlphaFold model of the $\sigma^A{}_4$-bound Q857R7 (pale green) with that of WhiB1 (PDB ID:6ONO, pink) at the Wbl:$\sigma^A{}_4$ interface around the [4Fe-4S] cluster binding pocket. The [4Fe-4S] cluster, the Cys ligands and the residues in the aromatic patch of WhiB1 are highlighted as the references. The Wbl residues corresponding to the aromatic patch in WhiB1 are highlighted in sticks with the C atoms colored pale green. Fe atoms are colored orange, S in yellow, O in red, and N in blue. $\sigma^A{}_4$ is not shown for clarity. The TM score of the AlphaFold model is 0.78 for Q857R7:$\sigma^A{}_4$. (B) and (C), UV-Visible spectra and SDS-PAGE analyses of the samples from co-expression and affinity purification of $\sigma^A{}_4$ and Q857R7 (wildtype and mutants as indicated) using Q857R7-His$_6$ as the bait. The absorption spectra were normalized to the protein concentration rather than to the absorbance at 280 nm due to the substantial differences in the 280-nm extinction coefficients between the Q857R7 proteins with and without binding to Strep-$\sigma^A{}_{C112}\beta_{tip}$. The absorption peak around 410 nm, highlighted with a dashed line, reflects the occupancy of the [4Fe-4S] cluster in the pull-down samples. The intensities of the 410-nm peak for the Q857R7 variant samples are comparable to each other, suggesting that the mutation of the aromatic residues does not disrupt the [4Fe-4S] cluster.

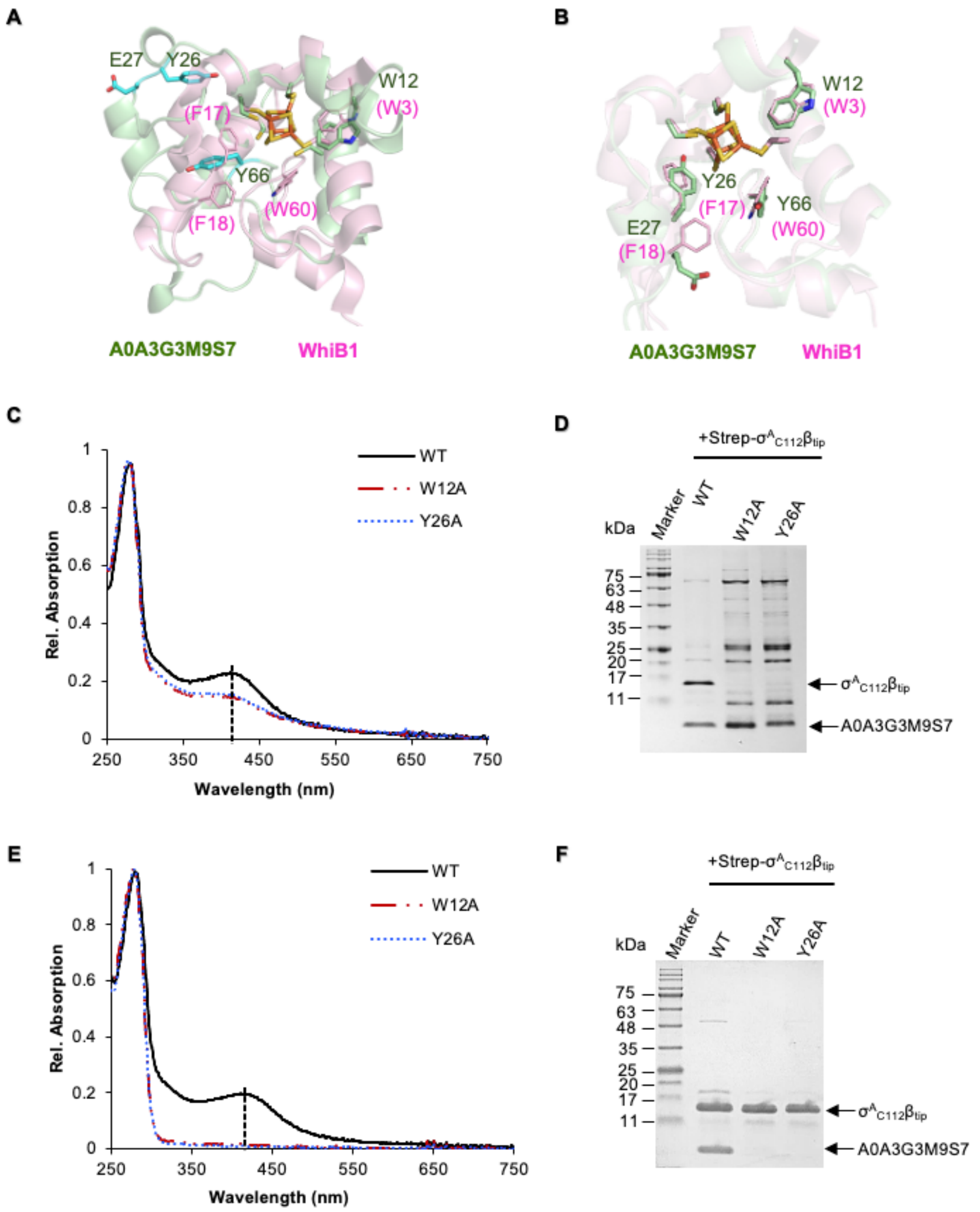


**Supplementary Figure 11. Effect of the aromatic patch residues in the Wbl (UniProt ID: A0A3G3M9S7) from *Gordonia phage Octobien14 Mycobacterium phage Lolly9* on $\sigma^A{}_4$ binding.** (A) and (B) Comparison of the AlphaFold model (Panel A) and SWISS-model (Panel B), respectively, of the $\sigma^A{}_4$-bound A0A3G3M9S7 (pale green) with that of WhiB1 (PDB ID:6ONO, pink) at the Wbl:$\sigma^A{}_4$ interaction interface around the [4Fe-4S] cluster binding pocket. The [4Fe-4S] cluster, the Cys ligands and the residues in the aromatic patch of WhiB1 are highlighted in sticks as the references. The Wbl residues corresponding to the aromatic patch residues in WhiB1 are highlighted in sticks with the C atoms colored pale green. The residues corresponding to the WhiB1 aromatic patch residue by sequence alignment but not by 3D structure alignment in Panel (A) are highlighted in sticks with the C atoms colored cyan. Fe atoms are colored orange, S in yellow, O in red, and N in blue. $\sigma^A{}_4$ is not shown for clarity. The TM score of the AlphaFold model is 0.64 for A0A3G3M9S7:$\sigma^A{}_4$. (C) and (D), UV-Visible spectra and SDS-PAGE analyses of the samples from co-expression and affinity purification of $\sigma^A{}_4$ and A0A3G3M9S7 (wildtype and mutants as indicated) using A0A3G3M9S7-$His_6$ as the bait. (E) and (F), UV-Visible spectra and SDS-PAGE analyses of the samples from co-expression and affinity purification of $\sigma^A{}_4$ and A0A3G3M9S7 (wildtype and mutants as indicated) using Strep-$\sigma^A{}_{C112}\beta_{tip}$ as the bait as an alternative strategy due to the high levels of contaminants shown in Panel (D). In all cases, the absorption spectra were normalized to the absorbance at 280 nm. The absorption peak around 410 nm, highlighted with a dashed line, reflects the occupancy of the [4Fe-4S] cluster in the pull-down samples. The lower intensities of the absorption peak at 410 nm for the W12A and Y26A mutant samples and higher levels of the contaminants in the pull-down samples (Panels C and D) compared to the wildtype suggest that these mutations destabilize the [4Fe-4S] cluster and overall folding of A0A3G3M9S7.

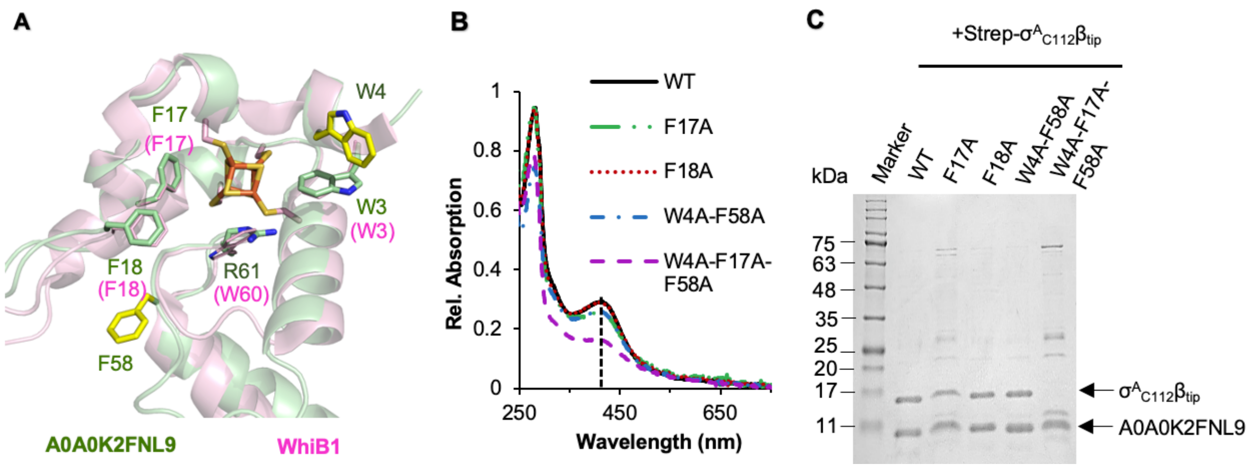


**Supplementary Figure 12. Effect of aromatic patch residues in the Wbl (UniProt ID: A0A0K2FNL9) from *Gordonia phage Octobien14* on $\sigma^A{}_4$ binding.** (A) Comparison of the AlphaFold model of the $\sigma^A{}_4$-bound A0A0K2FNL9 (pale green) with that of WhiB1 (PDB ID:6ONO, pink) at the Wbl:$\sigma^A{}_4$ interaction interface around the [4Fe-4S] cluster binding pocket. The [4Fe-4S] cluster, the Cys ligands and the residues in the aromatic patch of WhiB1 are highlighted in sticks as the references. The Wbl residues corresponding to the aromatic patch in WhiB1 are highlighted in sticks with the C atoms colored pale green. The two additional aromatic residues in A0A0K2FNL9 at the A0A0K2FNL9:$\sigma^A{}_4$. molecular interface but outside the [4Fe-4S] cluster binding site are highlighted in sticks with the C atoms in yellow. Fe atoms are colored orange, S in yellow, O in red, and N in blue. $\sigma^A{}_4$ is not shown for clarity. The TM score of the AlphaFold model is 0.66 for A0A0K2FNL9:$\sigma^A{}_4$. (B) and (C), UV-Visible spectra and SDS-PAGE analyses of the samples from co-expression and affinity purification of $\sigma^A{}_4$ and A0A0K2FNL9 (wildtype and mutants as indicated) using A0A0K2FNL9-His$_6$ as the bait. The absorption spectra of the mutants (W4A-F58A and W4A-F17A-F58A) were normalized to the protein concentration rather than to the absorbance at 280 nm due to the substantial differences in the 280-nm extinction coefficients between the A0A0K2FNL9 variants. The absorption peak around 410 nm, highlighted with a dashed line, reflects the occupancy of the [4Fe-4S] cluster in the pull-down samples. The intensities of the 410-nm peak for the A0A0K2FNL9 triple mutant sample, W4A-F17A-F58A, is significantly lower than the wildtype, suggesting that the tipple mutation destabilizes the [4Fe-4S] cluster.

### References


1. Wan, T., Li, S., Beltran, D. G., Schacht, A., Zhang, L., Becker, D. F., and Zhang, L. (2020) Structural basis of non-canonical transcriptional regulation by the $\sigma^A$-bound iron-sulfur protein WhiB1 in *M. tuberculosis*. *Nucleic Acids Res* **48**, 501-516
2. Wan, T., Horova, M., Beltran, D. G., Li, S., Wong, H. X., and Zhang, L. M. (2021) Structural insights into the functional divergence of WhiB-like proteins in *Mycobacterium tuberculosis*. *Mol Cell* **81**, 2887–2900
3. Shell, S. S., Wang, J., Lapierre, P., Mir, M., Chase, M. R., Pyle, M. M., Gawande, R., Ahmad, R., Sarracino, D. A., Ioerger, T. R., Fortune, S. M., Derbyshire, K. M., Wade, J. T., and Gray, T. A. (2015) Leaderless Transcripts and Small Proteins Are Common Features of the Mycobacterial Translational Landscape. *PLoS Genet* **11**, e1005641
4. Singh, A., Mai, D., Kumar, A., and Steyn, A. J. (2006) Dissecting virulence pathways of *Mycobacterium tuberculosis* through protein-protein association. *Proc Natl Acad Sci USA* **103**, 11346-11351
5. Wan, T., Horova, M., Khetrapal, V., Li, S., Jones, C., Schacht, A., Sun, X., and Zhang, L. (2023) Structural basis of DNA binding by the WhiB-like transcription factor WhiB3 in *Mycobacterium tuberculosis*. *The Journal of biological chemistry* **299**, 104777
6. Abramson, J., Adler, J., Dunger, J., Evans, R., Green, T., Pritzel, A., Ronneberger, O., Willmore, L., Ballard, A. J., Bambrick, J., Bodenstein, S. W., Evans, D. A., Hung, C. C., O'Neill, M., Reiman, D., Tunyasuvunakool, K., Wu, Z., Zemgulyte, A., Arvaniti, E., Beattie, C., Bertolli, O., Bridgland, A., Cherepanov, A., Congreve, M., Cowen-Rivers, A. I., Cowie, A., Figurnov, M., Fuchs, F. B., Gladman, H., Jain, R., Khan, Y. A., Low, C. M. R., Perlin, K., Potapenko, A., Savy, P., Singh, S., Stecula, A., Thillaisundaram, A., Tong, C., Yakneen, S., Zhong, E. D., Zielinski, M., Zidek, A., Bapst, V., Kohli, P., Jaderberg, M., Hassabis, D., and Jumper, J. M. (2024) Accurate structure prediction of biomolecular interactions with AlphaFold 3. *Nature* **630**, 493-500
